# Topology and contribution to the pore channel lining of plasma membrane-embedded *S. flexneri* type 3 secretion translocase IpaB

**DOI:** 10.1101/2021.10.07.463542

**Authors:** Poyin Chen, Brian C. Russo, Jeffrey K. Duncan-Lowey, Natasha Bitar, Keith Egger, Marcia B. Goldberg

## Abstract

*Shigella* spp. are human bacterial pathogens that cause bacillary dysentery.
Virulence depends on a type 3 secretion system (T3SS), a highly conserved structure present in multiple important human and plant pathogens. Upon host cell contact, the T3SS translocon is delivered to the host membrane, facilitates bacterial docking to the membrane, and enables delivery of effector proteins into the host cytosol. The *Shigella* translocon is composed of two proteins, IpaB and IpaC, which together form this multimeric structure within host plasma membranes. Upon interaction of IpaC with host intermediate filaments, the translocon undergoes a conformational change that allows for bacterial docking onto the translocon and, together with host actin polymerization, enables subsequent effector translocation through the translocon pore. To generate additional insights into the translocon, we mapped the topology of IpaB in plasma membrane-embedded pores using cysteine substitution mutagenesis coupled with site-directed labeling and proximity-enabled crosslinking by membrane permeant sulfhydryl reactants. We demonstrate that IpaB function is dependent on post translational modification by a plasmid-encoded acyl carrier protein. We show that the first transmembrane domain of IpaB lines the interior of the translocon pore channel such that the IpaB portion of the channel forms a funnel-like shape leading into the host cytosol. In addition, we identify regions of IpaB within its cytosolic domain that protrude into and are closely associated with the pore channel. Taken together, these results provide a framework for how IpaB is arranged within translocons natively delivered by *Shigella* during infection.

**Importance:** Type 3 secretion systems are nanomachines employed by many bacteria, including *Shigella*, which deliver into human cells bacterial virulence proteins that alter cellular function in ways that promote infection. Delivery of *Shigella* virulence proteins occurs through a pore formed in human cell membranes by the proteins IpaB and IpaC. Here, we define how IpaB contributes to the formation of pores natively delivered into human cell membranes by *S. flexneri*. We show that a specific domain of IpaB (transmembrane domain 1) lines much of the pore channel and that portions of IpaB that lie in the inside of the human cell loop back into and/or are closely associated with the pore channel. These findings provide new insights into the organization and function of the pore in serving as the conduit for delivery of virulence proteins into human cells during *Shigella* infection.

## Introduction

*Shigella* spp., Gram-negative bacterial gastrointestinal pathogens, are a main cause of childhood mortality worldwide (1). *Shigella* establish infection via a type 3 secretion system (T3SS), a nanomachine used by pathogens to inject effector proteins into the cytosol of host cells (2). The T3SS forms a translocon in the plasma membrane of host cells and delivers through the pore virulence proteins that facilitate bacterial uptake into host cells causing infection (3–7). The translocon enables bacterial docking to the cell and, in contrast to being a passive portal, actively participates in regulation of effector secretion into the cell (8). Specifically, emerging evidence indicates that the pore is actively involved in regulating bacterial docking and effector secretion (9, 10).

T3SS translocon are composed of two protein translocases - in *Shigella*, IpaB and IpaC - which are conserved among T3SS pathogens (5). IpaC is a single pass transmembrane protein with an N-terminal extracellular domain, cytosolic residues C-terminal to the transmembrane domain, and residues proximal to the C-terminus that reside within the translocon pore channel interior (11). Following membrane insertion, the interaction of C-terminal residues in IpaC with host intermediate filaments induces a conformational change in the translocon, allowing docking of the *Shigella* T3SS needle onto the pore (11). In conjunction with IpaC binding intermediate filaments, polymerization of host actin triggers opening of the pore channel, enabling the delivery of effector proteins into the host cytosol (9).

IpaB is a two-pass transmembrane protein (Fig. 1A), with both N and C-termini facing the extracellular space (2). Previous studies on IpaB largely focused on functional analyses and oligomeric behavior using purified and recombinant IpaB (12, 13). Targeted deletion of IpaB residues surrounding and encompassing the transmembrane domains (TM1 and TM2) negatively impact IpaC recruitment to host membranes (14), suggesting that IpaB interactions with IpaC involve the IpaB transmembrane domains. Studies using purified, recombinant IpaB show that in mild detergents, IpaB forms tetramers (12, 13), consistent with the model that translocons are hetero-oligomeric structures. Proteolysis studies with IpaB in liposomes defined the IpaB transmembrane domains to include amino acid residues 313-333 (transmembrane domain 1; TM1) and 400-419 (TM2) (2). The organization of these domains and their interactions with other IpaB and IpaC molecules in a translocon are unknown.

**Figure 1.**
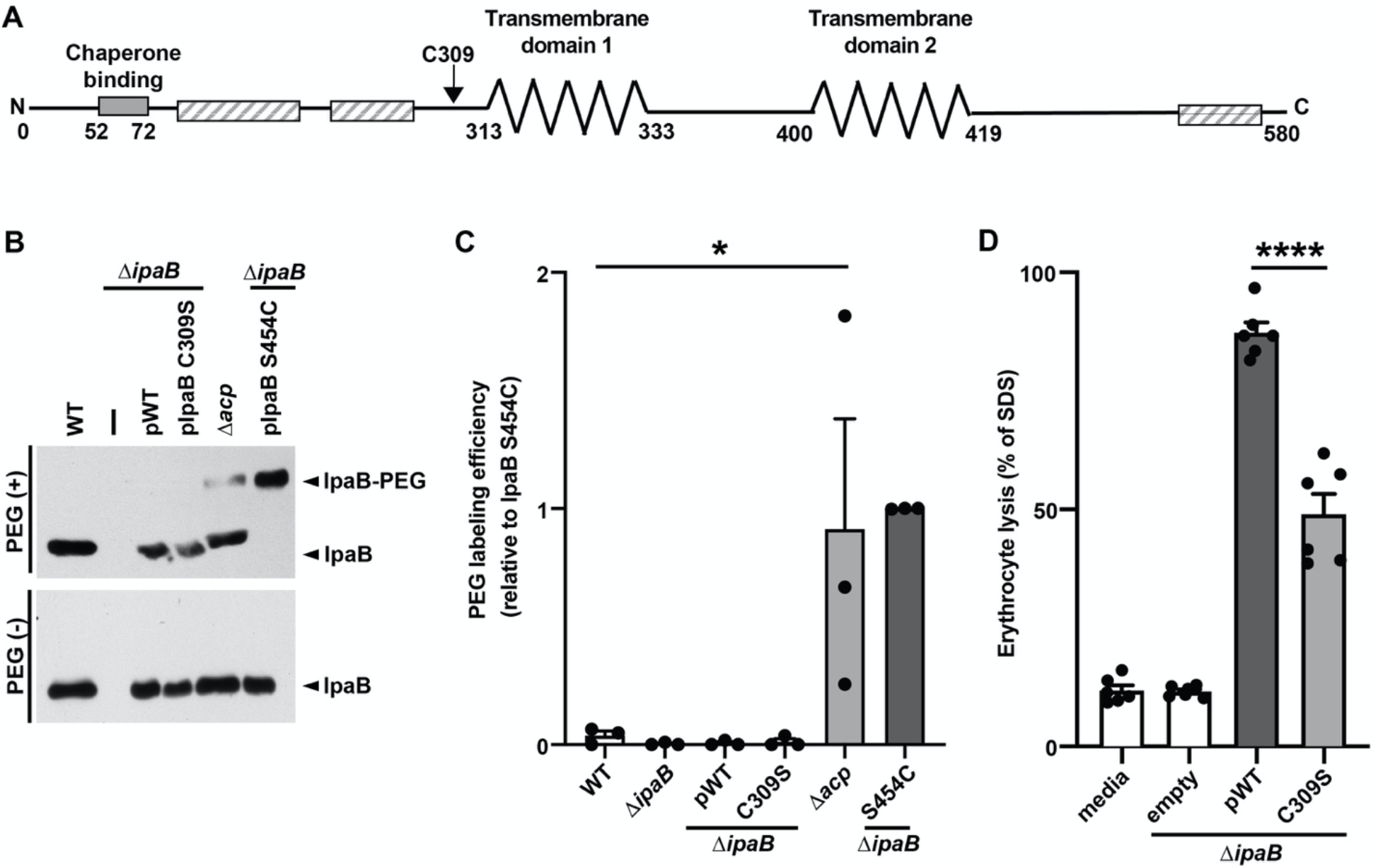
The IpaB native cysteine is sulfhydryl nonreactive and essential to translocon formation. (A) Schematic showing key features of IpaB. Bacterial chaperone binding site (solid gray bar), identified by experimental data. Transmembrane domains 1 (TM1) and 2 (TM2) (zigzag lines), predicted through *in silico* analyses and experimental data (2). Putative coiled-coiled regions (hatched bars), predicted by *in silico* analyses. The native cysteine, C309 (arrow). (B) Accessibility of the native cysteine in soluble IpaB. Gel migration of PEG5000-maleimide labeled (IpaB-PEG) or unlabeled (IpaB) soluble IpaB in culture supernatants of indicated strains following chemical activation of type 3 secretion with Congo red. Because expression of native IpaB is higher than of plasmid-borne IpaB, loading for WT and Δ*acp* (lanes 1 and 5) is 5-fold less than for all other strains. Wildtype *S. flexneri* (WT), *S. flexneri* Δ*ipaB*, *S. flexneri* Δ*ipaB* producing wildtype IpaB (pWT), IpaB C309S, or IpaB S454C, or *S. flexneri Dacp*. Representative western blot. (C) Densitometry of data represented in panel B. Plotted are means ± SEM. Black dots represent the values obtained from individual experiments. *, p<0.05; ANOVA with Dunnett’s *post hoc* test comparing IpaB labeling efficiency of *S. flexneri Dacp* to wildtype *S. flexneri*. (D) Quantification of *S. flexneri* pore formation by hemoglobin release following cocultured of indicated *S. flexneri* strains with sheep erythrocytes. *S. flexneri* expressing wildtype IpaB or *S. flexneri* Δ*ipaB* producing wildtype IpaB or IpaB C309S. Released hemoglobin was quantified at A_570_ from at least three independent experiments. Means ± SEM are plotted. Black dots represent the values obtained from individual experiments. ****, p<0.0001; ANOVA with Dunnett’s *post hoc* test comparing *S. flexneri* Δ*ipaB* producing IpaB C309S to *S. flexneri* Δ*ipaB* producing wildtype IpaB.

Structural investigations into the translocon, despite herculean efforts, have yet to define the structure of the pore with sufficient detail to determine its organization (15–17). Understanding how IpaB contributes to the organization and function of plasma membrane-embedded translocons is important to comprehensively understand the role of translocons in pathogenesis. Our studies here into the topology of IpaB provide additional insights into the arrangement of the translocases within natively delivered, functional translocons. Using cysteine accessibility mutagenesis, we achieve single-residue resolution of the topology of IpaB within membrane-embedded translocons. We show that TM1 of IpaB lines much of the interior of the pore channel, whereas aside from a single residue, TM2 is undetectable from the pore channel and therefore likely largely embedded in the lipid bilayer. We demonstrate that residues within the cytosolic domain of IpaB immediately N-terminal to TM2 loop back into the translocon pore channel. Together with additional analysis using proximity-enabled crosslinking, our findings indicate that the IpaB portion of the translocon pore channel adopts a funnel-like conformation, wherein it narrows towards the cytosolic side of the plasma membrane. Together with evidence indicating that IpaC forms the very extracellular end of the pore channel (11), our data define the topology of the *Shigella* translocon and infer an overall shape of the pore.

## Results

### Approach for mapping the topology of IpaB

We mapped the topology of IpaB in membrane-embedded, natively delivered translocons using cysteine accessibility mutagenesis, replacing individual IpaB residues with cysteine and testing reactivity with sulfhydryl specific probes (Figs. 2A and 3A). Free (i.e., in a reduced state) and accessible sulfhydryl groups of cysteine residues will form a covalent bond with the probe. IpaB topology was defined by analyzing the accessibility of each substituted cysteine of IpaB in plasma membranes from HeLa cells infected with *S. flexneri* expressing individual cysteine derivatives in the presence of the sulfhydryl probe methoxypolyethylene glycol-maleimide (PEG5000-maleimide) (Fig. 3A). To control for inherent accessibility of the cysteine in the context of soluble protein, accessibility was also assessed in IpaB in culture supernatants following chemical activation of the T3SS (Fig. 2A).

**Figure 2.**
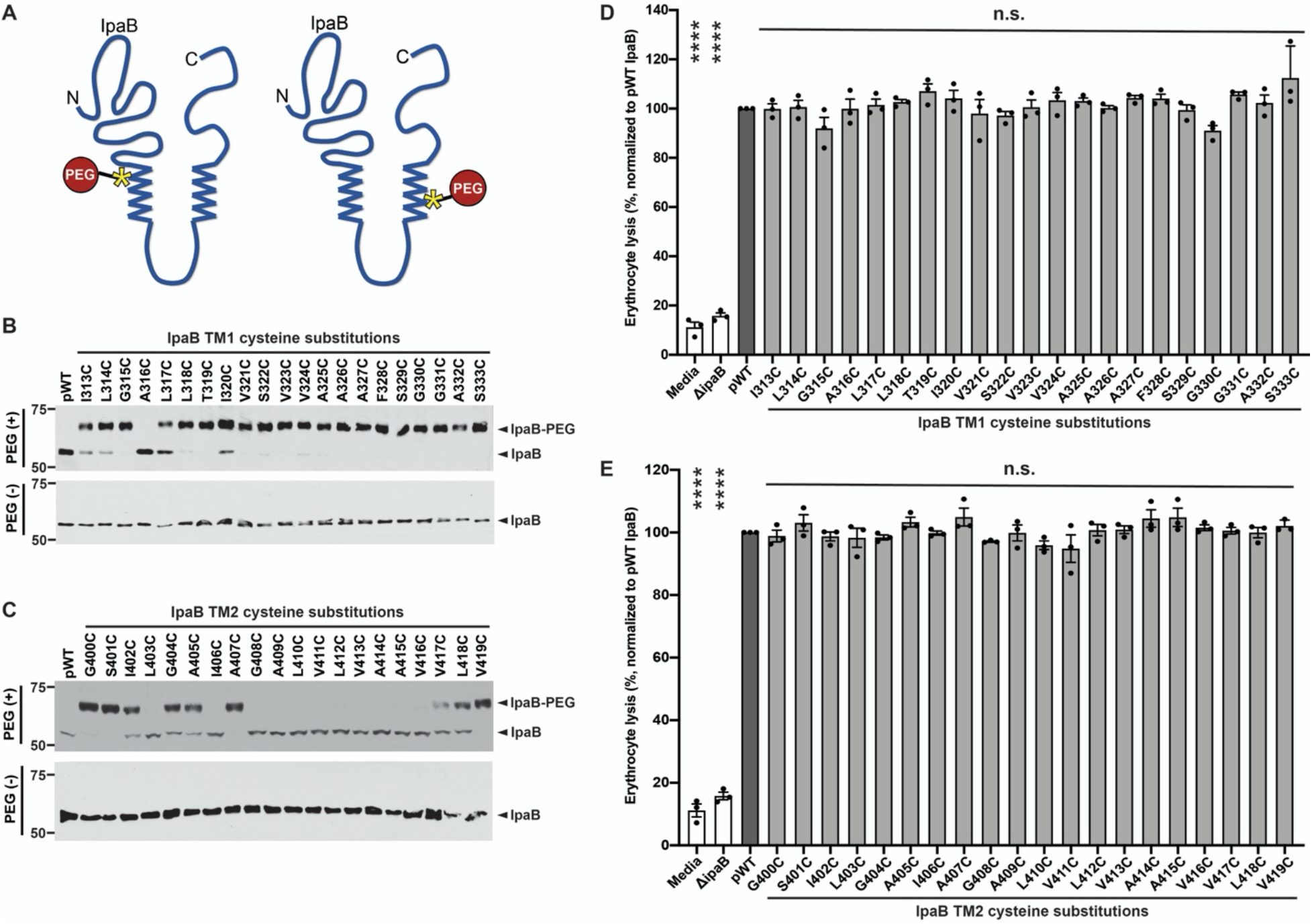
Cysteine substitutions along IpaB are variably accessible in the context of soluble IpaB and do not impair pore formation in mammalian membranes. (A) Schematic showing PEG5000-maleimide labeling of soluble *S. flexneri* IpaB cysteine substitution derivatives. Transmembrane domains (zigzag lines), representative cysteine substitution derivatives along the length of IpaB (asterisks), and PEG5000-maleimide (red circle) are depicted. (B and C) Gel migration of PEG5000-maleimide labeled (IpaB-PEG) or unlabeled (IpaB) soluble IpaB in culture supernatants of indicated strains following chemical activation of type 3 secretion with Congo red.*S. flexneri* Δ*ipaB* expressing wildtype IpaB (pWT) or a single IpaB cysteine substitution derivative. Representative western blots. (D and E) Quantification of hemoglobin release by *S. flexneri* Δ*ipaB* expressing wildtype IpaB (pWT) or a single cysteine substitution derivative. The abundance of hemoglobin release was quantified at A_570_ from at least three independent experiments. Means ± SEM are plotted. Black dots represent values obtained from individual experiments. ****, p<0.0001; n.s., not significant; ANOVA with Dunnett’s *post hoc* test comparing each cysteine substitution mutant to *S. flexneri* Δ*ipaB* producing WT IpaB (pWT).

**Figure 3.**
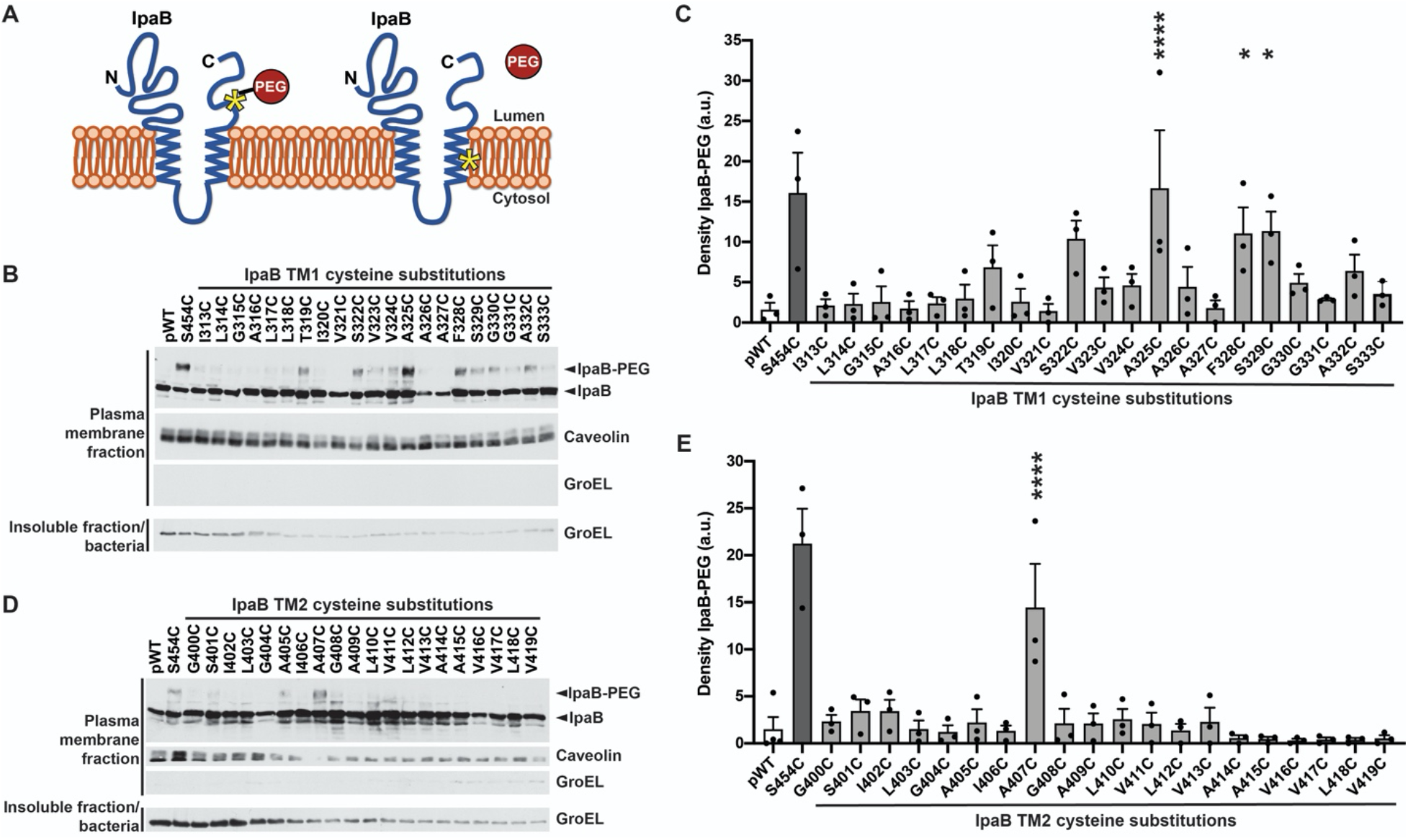
Cysteine accessibility of transmembrane domains in membrane embedded IpaB. (A) Schematic of IpaB topology mapping by PEG5000-maleimide labeling. The N and C-terminal domains of IpaB are situated on the extracellular side of the plasma membrane. Cysteine substitutions are represented by an asterisk. PEG5000-maleimide is membrane impermeant and size excluded from passing all the way through the translocon pore, therefore does not react with cysteines on the cytosolic side of the plasma membrane nor with cysteines facing the lipid bilayer. (B and D) Gel migration of PEG5000-maleimide labeled (IpaB-PEG) or unlabeled (IpaB) IpaB in membrane inserted translocons. *S. flexneri* Δ*ipaB* expressing wildtype IpaB (pWT) or a single IpaB cysteine substitution derivative. Positive control, IpaB S454 lies in the C-terminal extracellular domain; S454C is accessible to PEG5000-maleimide labeling. Representative western blots are shown. Caveolin-1, plasma membrane marker; GroEL, bacterial cytosolic protein. (C and E) Densitometry analysis of IpaB-PEG5000 bands from three independent experiments represented in panels B and D. Means ± SEM are plotted. Black dots represent values obtained from individual experiments. *, p<0.05; ****, p<0.0001; ANOVA with Dunnett’s *post hoc* test comparing IpaB labeling efficiency of *S. flexneri* Δ*ipaB* expressing each IpaB cysteine derivative to *S. flexneri* Δ*ipaB* expressing WT IpaB (pWT); the difference for each other strain is not significant.

We selected PEG5000-maleimide, the approximate diameter of which is 4.4 nm (11), because it is too large to pass through the pore, which is predicted to be 2.5 nm at its narrowest point (5). By adding PEG5000-maleimide to the extracellular media during infection, our approach assesses the accessibility of IpaB residues specifically from the exterior of the cell (11, 18).

### The IpaB native cysteine (C309) is post-translationally acylated by virulence plasmid-encoded *acp* and is required for translocon function

Reactive sulfhydryl groups present in the native protein being analyzed confound labeling results. IpaB contains a single native cysteine at residue 309, located in the N-terminal extracellular domain immediately N-terminal to the start of the first transmembrane domain (TM1; residues 313-333) (Fig. 1A). To determine whether IpaB C309 is free and accessible in soluble, secreted IpaB (as opposed to membrane embedded IpaB), we tested the reactivity of IpaB C309 with PEG5000-maleimide in bacterial culture supernatants obtained under T3SS activating conditions.

PEG5000-maleimide labeling of IpaB in culture supernatants from wildtype bacteria or an *ipaB* mutant producing either wildtype IpaB or an IpaB derivative lacking the native cysteine (IpaB C309S) showed no detectable size shift corresponding to cysteine reactivity to maleimide (Fig. 1B), whereas the control cysteine substitution IpaB S454C, located in the IpaB C-terminal extracellular domain, showed a band that migrated at the expected size of IpaB labeled with PEG5000-maleimide (Fig. 1B, right-most lane). The absence of significant labeling for wildtype IpaB (Fig. 1B and C) indicates that the native cysteine of IpaB is inaccessible to PEG5000-maleimide.

The *Salmonella* homolog of IpaB, SipB, contains a cysteine that is conserved with IpaB C309 and is acylated in the bacterial cytoplasm by an acyl carrier protein (19). Since *S. flexneri* also encodes on its virulence plasmid an acyl carrier protein (Acp), we postulated that IpaB C309 is acylated by Acp. Deletion of the *acp* enabled partial sulfhydryl reactivity of IpaB C309 (Fig. 1B and C), suggesting that IpaB is post translationally modified on residue C309 by Acp. Site-directed mutagenesis of IpaB C309 to a serine (C309S) resulted in a significant decrease in pore formation (Fig. 1D), as measured by the ability to lyse sheep erythrocytes, indicating that the cysteine at position 309 of IpaB is essential for formation of translocon pores. Thus, IpaB C309 is prevented from reacting with PEG5000-maleimide by Acp modification and analysis of cysteine substitution mutants can be performed with the IpaB native cysteine at position 309, which is essential for the function of the *S. flexneri* translocon.

### Cysteine substitutions at multiple positions are accessible to PEG5000-maleimide and are tolerated for IpaB function

IpaB residues in TM1, the cytosolic domain, the second transmembrane domain (TM2) (2), and the C-terminal extracellular domain (Fig. 1A) were replaced with cysteine and introduced into the *ipaB* mutant. Resulting strains were each assessed for host invasion (Fig. S1) and accessibility of the substituted cysteine residue in the context of secreted, soluble IpaB (Fig. 2A and S2). In soluble protein, all cysteine substitutions within TM1 except IpaB A316C and most in the cytosolic domain reacted with PEG5000-maleimide (Figs. 2B and S2). In contrast, many cysteine substitutions within IpaB TM2 (IpaB L403C, I406C, and G408C-V416C) showed no detectable PEG5000-maleimide reactivity (Fig. 2C). Although we cannot definitively exclude the possibility that the absence of TM2 cysteine labeling is due to aberrant acylation of the substituted cysteine, this seems unlikely because the acylated cysteine of SipB and the putatively-acylated cysteine of IpaB are within a conserved sequence motif associated with acylation (Met-Gly-Cys-Val/Ile-Gly-Lys-Ile/Val, (19)) that is absent at the TM2 cysteine substitutions. That this conserved sequence constitutes an acyl carrier recognition motif is strongly suggested by its conservation in SipB homologs that are genetically linked with acyl carrier genes but not in SipB homologs not linked with acyl carrier genes (19). The lack of cysteine labeling was not due to a lack of protein, since similar quantities of IpaB were detected for all IpaB mutants (Figs. 2B-C and S2).

Cysteine substitution derivatives were tested for their ability to support translocon pore formation, by assaying hemolysis of sheep erythrocytes. Each cysteine substitution derivative in TM1 and TM2 and all but four substitutions in the cytosolic and extracellular domains (IpaB D347C, S379C, A381C, and A431C) supported the formation of translocon pores at levels similar to that of wildtype IpaB (Figs. 2D and E and S3). Together, these data show that cysteine incorporation at any of numerous IpaB residues does not alter IpaB function, as pore formation (Figs. 2 and S3) and invasion (Fig. S1) occur at rates similar to WT IpaB. Only those cysteine substitution derivatives that supported pore formation and invasion at an efficiency comparable to wildtype IpaB were included in subsequent studies.

### TM1 of IpaB lines much of the interior of the translocon pore channel

The topology of membrane-embedded IpaB in a natively delivered translocons was determined by assessing the accessibility of IpaB residues from the extracellular side of the membrane using PEG5000-maleimide labeling during *S. flexneri* infection of HeLa monolayers (Fig. 3A). Within TM1, IpaB S322C, A325C, F328C, and S329C showed an additional band that migrated more slowly by SDS-PAGE, consistent with PEG5000-maleimide labeling (Fig. 3B). Additional residues, predominantly within the C-terminal half of TM1, displayed labeling that upon densitometry did not reach statistical significance (Fig. 3B-C). In contrast, wildtype IpaB showed significantly less density at a similar position on the blot (Fig. 3C; p<0.1 for IpaB S322C; p<0.05 for IpaB A325C, F328C, and S329C). As observed here, for proteins embedded in plasma membranes, labeling of residues is typically only partial (11).

In contrast to IpaB TM1, the only cysteine substitution derivative in IpaB TM2 to show significant PEG5000-maleimide labeling was IpaB A407C (Fig. 3D and E; p<0.05). With only a single residue in IpaB TM2 that is accessible to sulfhydryl reactivity when in the membrane-embedded translocon, it is likely that in pores, the bulk of TM2 is positioned in the membrane away from the pore channel such that IpaB TM2 is embedded within the lipid bilayer and/or interacts with other IpaB and/or IpaC transmembrane regions such that PEG5000-maleimide accessibility of TM2 residues is blocked.

The accessibility to PEG5000-maleimide labeling of multiple residues within IpaB TM1 (Fig. 3B and C), the accessibility of only one residue within IpaB TM2 (Fig. 3C and D), and the previous observation that only one residue within the single TM of IpaC is accessible (11) together suggest that TM1 of IpaB constitutes the lining of much of the translocon pore channel.

### In adjacent IpaB molecules within intact translocons, residues near the cytosolic end of IpaB TM1 are close to one another

To identify individual IpaB residues that lie immediately adjacent to one another in neighboring IpaB molecules within translocons, we used proximity-triggered covalent crosslinking of IpaB cysteine derivatives induced by the presence of the membrane-permeant oxidant copper phenanthroline. Crosslinking of IpaB molecules requires that their cysteine residues are close enough to one another to form a disulfide bond, the length of which is estimated to be 2.05 Å (20), are in an orientation that enables the bond to form, and are located in a non-reducing environment. Since the host cell cytosol is reducing, residues that form disulfide bonds in response to oxidants are located either in the extracellular domain or within the pore channel.

Addition of copper phenanthroline to the infection medium resulted in the appearance of slower migrating IpaB-specific bands at a molecular weight consistent with the formation of IpaB dimers. The formation of IpaB dimers was consistently most prominent in a stretch of cysteine residues in IpaB TM1 that lie in the cytosolic half of the membrane-embedded alpha helix (IpaB S322C, F328C-S333C; Fig. 4). As observed here, crosslinking of residues in membrane-embedded translocons is typically incomplete (11).

**Figure 4.**
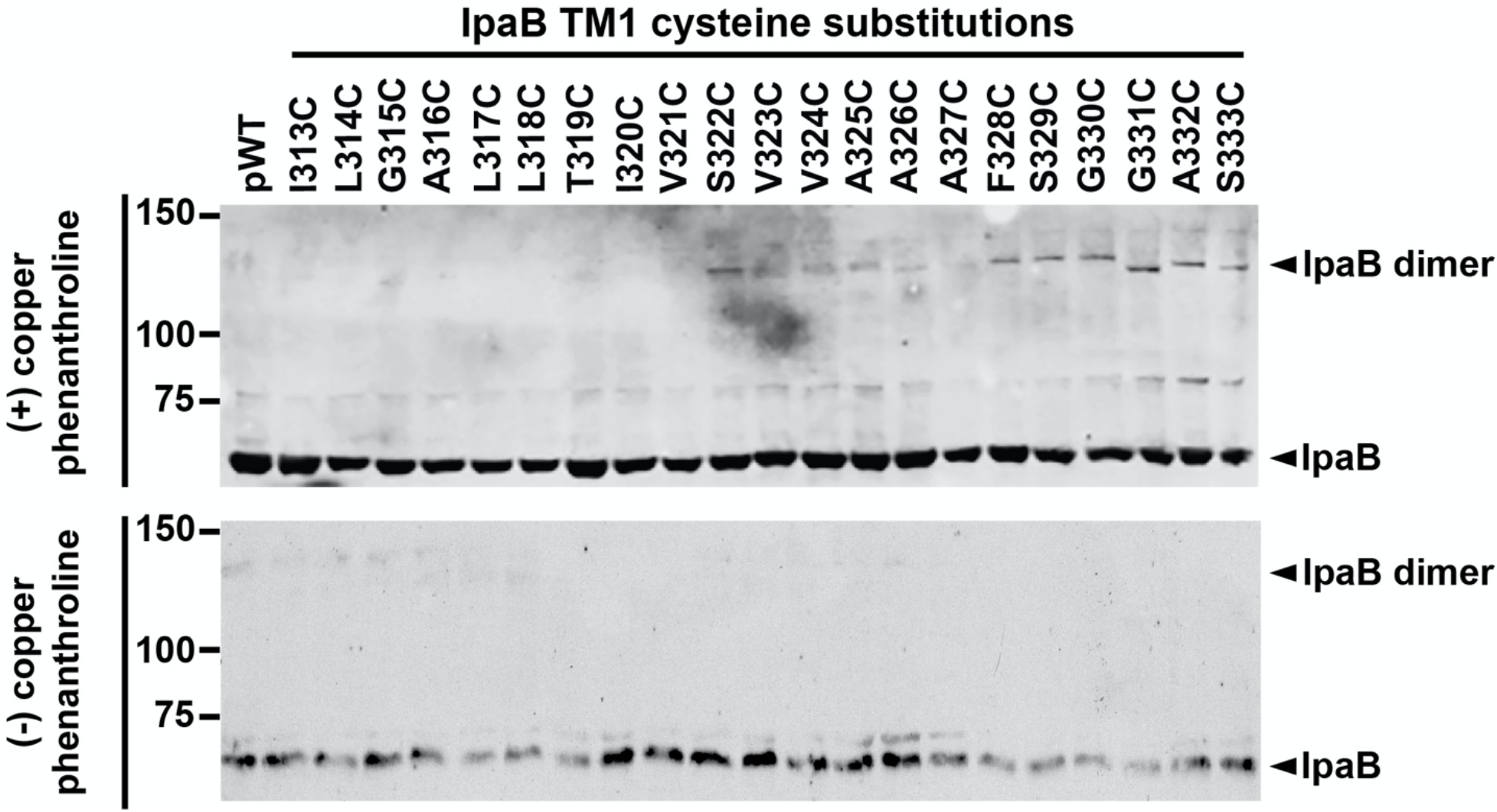
Proximity-enabled copper phenanthroline crosslinking of IpaB TM1 residues. Gel shift of copper 1,10-phenanthroline crosslinking of membrane embedded IpaB. Representative western blots are shown. The positions of IpaB (IpaB) and crosslinked IpaB (IpaB dimer) are shown to the right of the gel. Size markers are shown to the left of the gel.

The dimerization of residues in IpaB TM1 that lie close to the cytosolic side of the membrane (Fig. 4) are consistent with a model in which the portion of the translocon pore channel approaching the cytoplasm, which is made up of TM1, narrows in such a way that adjacent IpaB molecules are sufficiently close to one another to form crosslinks. No IpaB dimers were detected for TM2 cysteine substitutions, indicating that when organized in a membrane-embedded translocon, TM2 residues in adjacent IpaB molecules are insufficiently close to one another to crosslink in the presence of oxidant and/or are facing the bilayer such that they are inaccessible to membrane-permeant small molecules.

### The cytosolic domain of IpaB loops into the pore channel interior

Topological studies of the second translocase IpaC demonstrated that in membrane-embedded pores a stretch of 15 residues within the IpaC cytosolic domain loops back into the channel of the translocon pore (11). To assess whether residues within the cytosolic domain of IpaB also loop back into the translocon pore channel, the accessibility from the extracellular side of cells of a subset of residues within the IpaB cytosolic domain was assessed with PEG5000-maleimide labeling. HeLa cells were infected with *S. flexneri* strains expressing single cysteine substitution derivatives in this domain. IpaB A369C, E387C, G388C, G390C, D392C-K395C showed PEG5000 labeling that was significant (Fig. 5A and 5B). Among these residues, oxidant-induced dimerization was most prominent for K395C (Fig. 5C), indicating that K395 residues in neighboring IpaB molecules are in close proximity. Thus, the IpaB cytosolic domain, particularly those cytosolic residues that lie immediately N-terminal to TM2, are accessible from the extracellular surface of the plasma membrane. These results indicate that as for a stretch of residues in the cytosolic domain of IpaC (11), a stretch of cytosolic IpaB residues loops back into the channel of the translocon pore. Moreover, IpaB K395C sulfhydryl reactivity in both an extracellular accessibility (PEG5000-maleimide) and a proximity-enabled (copper phenanthroline) manner that is limited to non-reducing environments indicates that this residue clearly lies within the channel of the translocon pore; as it is five residues N-terminal to the start of IpaB TM2, it localizes to the portion of the pore channel near the cytoplasm.

**Figure 5.**
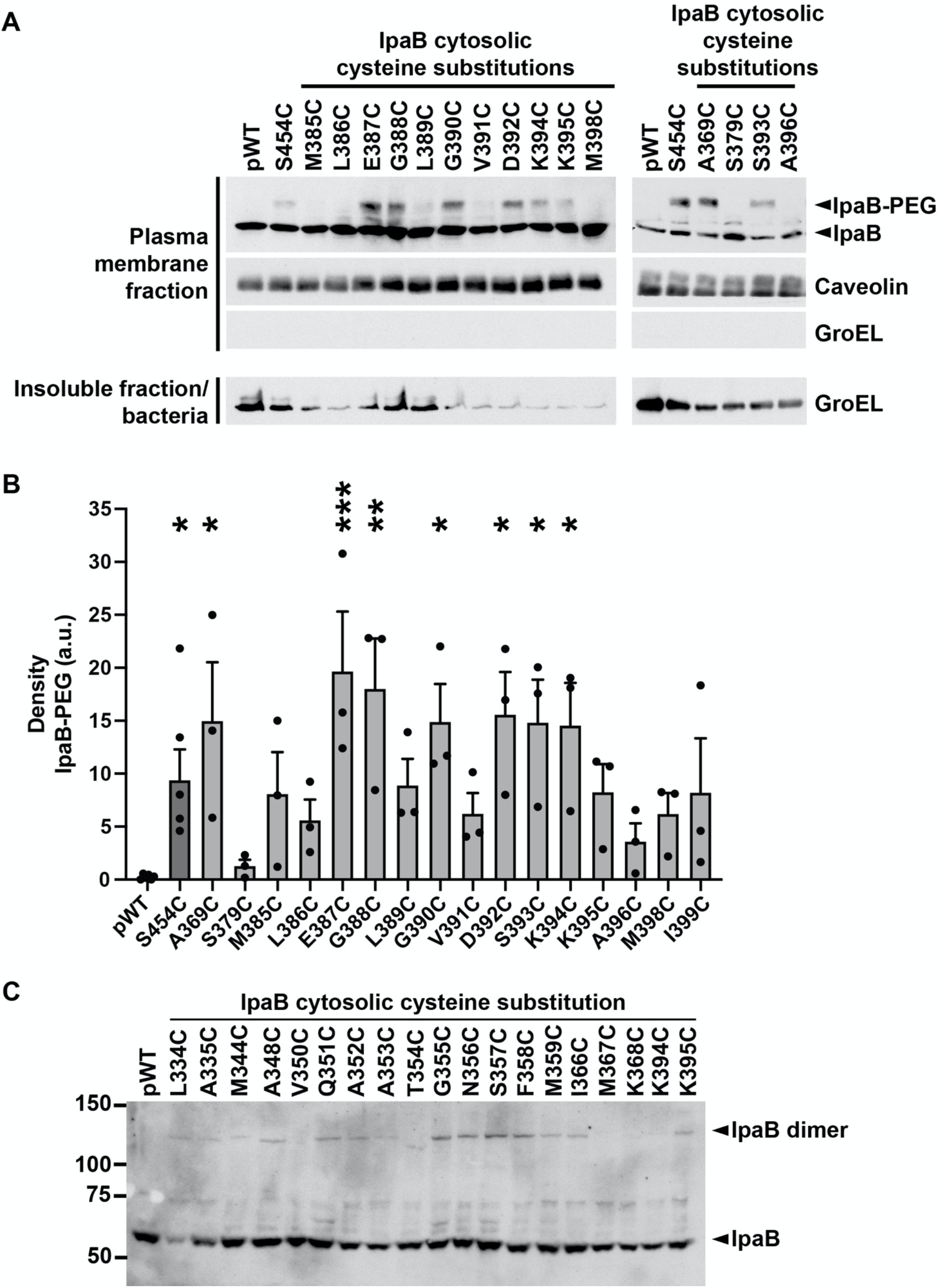
When in membrane embedded translocons, the cytosolic domain of IpaB is accessible to PEG5000 from the extracellular side of the plasma membrane. (A) Gel shift of PEG5000-maleimide labeled, membrane embedded IpaB. *S. flexneri* Δ*ipaB* expressing wildtype IpaB (pWT) or a single IpaB cysteine substitution derivative. Positive control, IpaB S454 lies in the C-terminal extracellular domain; S454C is accessible to PEG5000-maleimide labeling. Representative western blots. The positions of PEG5000-maleimide labeled IpaB (IpaB-PEG), unlabeled IpaB (IpaB), caveolin-1, a eukaryotic membrane marker, and GroEL, a bacterial cytosolic protein, are shown to the right of the gel. (B) Densitometry analysis of IpaB-PEG5000 bands from three independent experiments, represented in panel A. Means ± SEM are plotted. Black dots represent values obtained from individual experiments. *, p<0.05; **, p<0.01; ***, p<0.001; ANOVA with Dunnett’s *post hoc* test comparing IpaB labeling efficiency of *S. flexneri* Δ*ipaB* expressing each IpaB cysteine derivative to *S. flexneri* Δ*ipaB* expressing WT IpaB (pWT); the difference for each other strain is not significant. (C) Gel shift of copper 1,10-phenanthroline crosslinking of membrane inserted IpaB. Representative western blots are shown. The positions of IpaB (IpaB) and crosslinked IpaB (IpaB dimer) are shown to the right of the gel. Size markers are shown to the left of the gel.

## Discussion

In all T3SS-encoding bacteria, the translocon is an essential component of the T3SS, without which delivery of bacterial effectors into the host cell does not occur. Our understanding of the role of T3SS translocons has evolved from a passive opening for delivery of bacterial virulence proteins into the host cell cytosol to a pore that regulates T3SS activity. To generate insights into how the translocon is involved in the regulation of effector secretion, we mapped the topology of IpaB in a natively delivered, plasma membrane-embedded pore.

We show that the conserved native cysteine at position 309 of IpaB is essential for formation of translocons in membranes and that this cysteine is not accessible to sulfhydryl reactants such as PEG5000-maleimide (Fig. 1). This lack of sulfhydryl reactivity is likely due to posttranslational acylation of the cysteine by the virulence plasmid-encoded acyl carrier protein, Acp, as deletion of *acp* enables partial reactivity of C309 to PEG5000-maleimide. These results are consistent with findings reported for SipB, the *Salmonella* homolog to IpaB, in which the SipB conserved cysteine (C316) is post translationally acylated by the acyl carrier protein IacP, a homolog of *Shigella* Acp, and mutation of the native cysteine to alanine significantly decreased both the insertion of SipB into erythrocyte membranes and *Salmonella* entry into host monolayers (19). In both systems, mutation of the native cysteine resulted in a 40% decrease in the efficiency of pore formation (Fig. 1D and (19), respectively) suggesting this modification is generally required for translocon function. In contrast to the findings for SipB, deletion of plasmid-encoded *acp* in *S. flexneri* does not enable fully sulfhydryl reactivity of IpaB C309 (Fig. 1B and C). This discrepancy could be attributed to acyl carrier protein activity from the copy of *acp* that is present on the *Shigella* chromosome, steric hinderance of IpaB C309 due to IpaB conformation, and/or differences in methodology. The location of IpaB C309 immediately N-terminal to the first transmembrane domain, suggests that in functional translocons, it lies directly against the plasma membrane of the host cell and that the presumed addition of an acyl group to this residue contributes to efficient targeting and/or insertion of IpaB into membranes during translocon formation.

Transmembrane domains are alpha helices with approximately 3.6 amino acids per turn (21). TM1 and TM2 of IpaB are each predicted to be 20 amino acids (2), suggesting that that each IpaB transmembrane domain alpha helix contains five to six turns. Beginning with IpaB T319C, cysteine substitution mutations along IpaB TM1 showed reactivity to PEG5000-maleimide labeling that trended toward significant (or was significant) every 3-4 residues, approximating every full turn (Fig. 3B and C). In contrast to the observed periodic spacing of residue accessibility to PEG5000-maelimide, TM1 residues F328-S333, which lie closest to the cytosolic side of the bilayer, all crosslink efficiently upon exposure to oxidant, suggesting that the conformational changes that occur upon bacterial docking and effector translocation are associated with rotational flexibility in the TM1 alpha helix. These patterns of labeling indicate that the first transmembrane domain of IpaB lines much of the translocon channel.

In contrast to TM1, when in a membrane-embedded translocon, IpaB TM2 is largely inaccessible to PEG5000-maleimide labeling (Fig. 3D and E). The lack of detectable PEG5000-maleimide labeling in membrane-embedded IpaB L403C, I406C, and G408C-V416C was unsurprising given the lack of accessibility of these residues in soluble IpaB (Fig. 2C). Their inaccessibility in soluble IpaB was not due to interaction with IpaC, since when secreted from an *ipaC* mutant, they were inaccessible, but could be attributed to intramolecular steric hinderance from IpaB folding or oligomerization (12, 13) and/or to interactions of this stretch of IpaB with another protein that is established upon secretion of IpaB and prior to IpaB insertion into the membrane.

Conversely, IpaB cysteine derivatives at TM2 residues G400C-I402C, G404C, A405C, and V417C-V419C demonstrated efficient PEG5000-maleimide labeling in soluble IpaB yet PEG5000-maleimide inaccessibility in membrane-embedded IpaB (Figs. 2C and 3C). Given our evidence that IpaB TM1 lines much of the pore channel, the absence of accessibility of these TM2 residues is most likely because these portions of the TM2 alpha helix face the lipid bilayer. Alternatively, they could be involved in IpaB interactions established upon membrane insertion.

In membrane-embedded IpaB, but not in soluble secreted IpaB, for each residue that labeled with PEG5000-maleimide, labeling was incomplete, in that a subpopulation of IpaB molecules did not react with the PEG5000-maleimide (Fig. 3), as was previously observed for IpaC (11). A likely explanation for incomplete labeling of any cysteine in the translocon is that the presence of a PEG5000 adduct sterically blocks accessibility of additional PEG5000 molecules. In the absence of IpaC, IpaB inserts into membranes (2, 9, 22), raising the possibility that during bacterial infection, some IpaB inserts in the membrane separate from translocons; thus, also contributing to incomplete labeling could be membrane associated IpaB that is not in translocons and which does not label. Supporting this possibility, in the absence of *ipaC*, membrane delivered IpaB S454C is unlabeled (Fig. S4), a finding that also suggests that the conformation of membrane-associated translocon-independent IpaB differs from that of IpaB in translocons.

Only a single residue of the IpaC transmembrane domain, A106C, is accessible to extracellular PEG5000-maleimide labeling in membrane-embedded pores (11), which together with the data presented here, indicates that, like IpaB TM2, most of the IpaC transmembrane domain is likely facing the lipid bilayer. IpaC A106 is located close to the extracellular face of the translocon, the portion of the channel for which IpaB TM residues are poorly accessible.

We determined the proximity of an IpaB cysteine residue in one IpaB molecule to the same residue in neighboring IpaB molecules within intact membrane-embedded translocons using proximity-enabled disulfide bond crosslinking with the membrane permeant oxidant copper phenanthroline. Since disulfide bonds have a length of 2.05Å (20), dimerization of IpaB cysteine substitution mutants indicates that these specific sites on IpaB are within this distance of one another. Consistent with atomic force microscopy studies demonstrating that the T3SS translocon pore is funnel shaped (16), our data show that cysteine residues in adjacent IpaB molecules in the portion of the pore channel closest to the cytosol form crosslinks (IpaB TM1 cysteine derivatives S322C and F328C-S333C, Fig. 4), indicating that this part of the channel is narrow, whereas TM1 residues toward the extracellular face of the membrane are farther apart. The absence of complete dimerization of specific IpaB residues could be due to steric hinderance against additional covalent crosslinking and/or pore deformation from the initial crosslinking, causing other IpaB molecules in the pore to move farther away from one another, no longer permitting crosslinking.

Based on PEG5000-maleimide accessibility, IpaB TM1 residues S322, F328, and S329 line the pore channel (Fig. 3B), and based on copper crosslinking, are close to the same residue in adjacent IpaB molecules (Fig. 4). IpaB V324, A325, and A326, which also line the pore channel, only weakly crosslinked, suggesting that, in adjacent IpaB molecules, they are farther apart and consistent with the IpaB component of pore channel being wider towards the extracellular face of the plasma membrane (Fig. 6).

**Figure 6.**
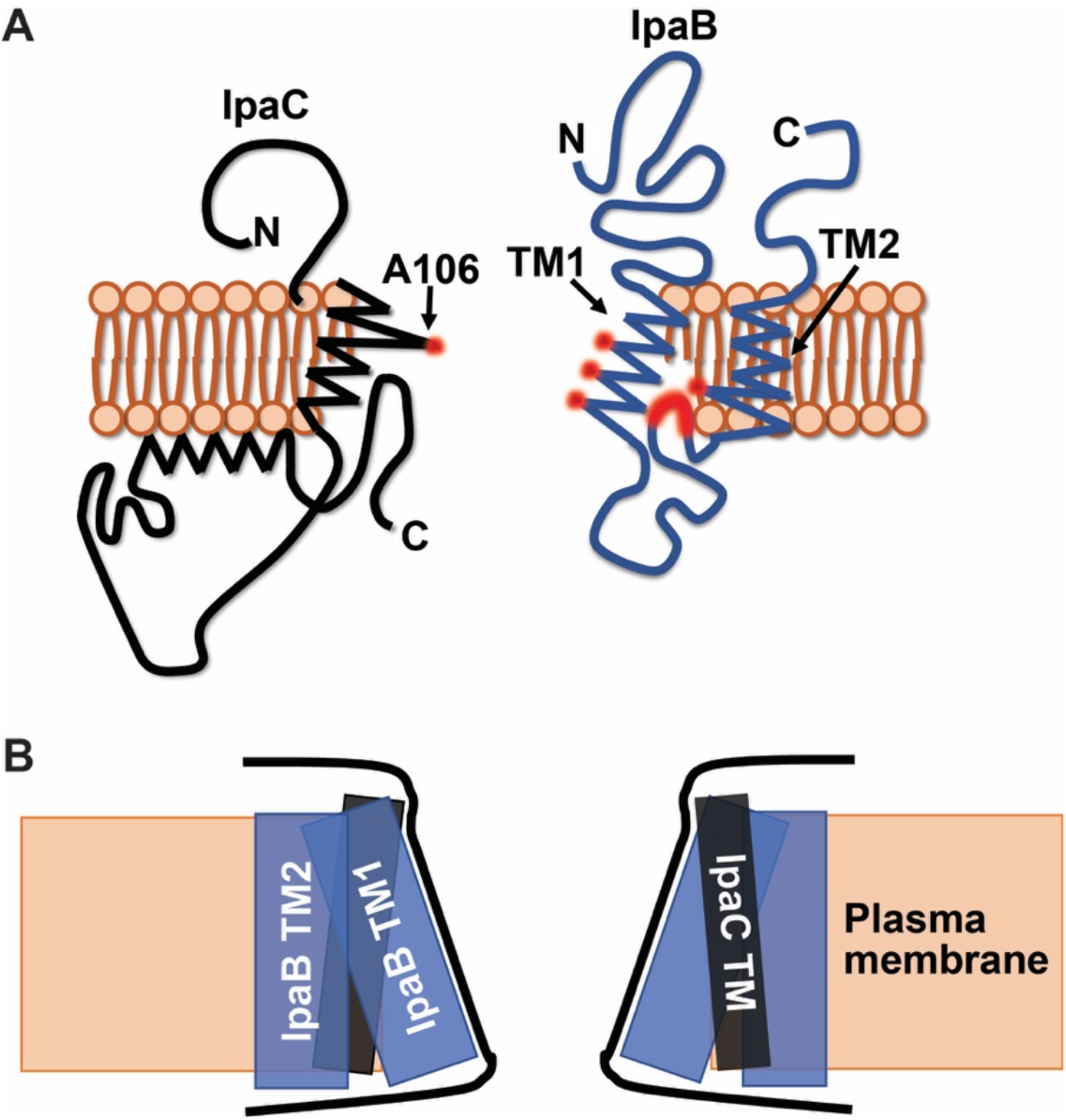
Model of IpaB and IpaC in membrane-embedded translocons. (A) Schematic of IpaB (blue) and IpaC (black) in the plasma membrane (orange). Residues in the transmembrane domains that are extracellularly accessible to PEG5000-maleimide highlighted in red. Whereas shown is only one molecule of each translocase, the pore is hetero-oligomeric. Images are not drawn to scale. (B) Schematic of the shape of the channel of the *S. flexneri* translocon pore, formed by the transmembrane domains of IpaB and IpaC. IpaC TM (black rectangle), IpaB TM1 and TM2 (blue rectangles), plasma membrane (orange box). Proposed shape of the pore channel is outlined (black lines).

Within the transmembrane domain of IpaC, A106, which is near the extracellular face of the plasma membrane, is the only residue for which the cysteine derivative was accessible (11). We therefore propose that the portion of the IpaB transmembrane span that is close to the extracellular face of the bilayer is interspersed with the transmembrane domain of IpaC (Fig. 6).

We propose that the IpaB component of the translocon pore channel is shaped as a funnel where the spacing is wider at the extracellular end of the transmembrane domain and narrows approaching the cytoplasm. We propose that IpaB TM1 lines the interior of the translocon pore channel, IpaB TM2 faces the lipid bilayer, and the cytosolic residues immediately N-terminal to TM2 loop back into the pore channel (Fig. 6A). Together with our previous data on the topology of IpaC, the data presented here are consistent with IpaC constituting the extreme extracellular end of the pore channel lining and IpaB TM1 constituting the remainder of the lining of the pore channel (Fig. 6). The data demonstrating that IpaC A106C, which is adjacent to the extracellular face of the bilayer, and IpaB TM1 residues near the cytosolic face of the bilayer crosslink upon addition of copper oxidant (Fig. 4 and (11)) suggest that the pore tapers at each end and is widest from the middle to just before the extracellular face of the bilayer (Fig. 6B).

## Materials and Methods

### Bacterial strains and plasmids

The bacterial strains and plasmids used in this study are listed in Table S1. Wildtype *S. flexneri* is serotype 2a strain 2457T (LaBrec 1964). All mutants are isogenic derivatives of 2457T. Δ*ipaB* and Δ*acp* mutants were generated as described (23). *ipaB* derivatives were inserted into pDSW206; expression was induced with 1mM isopropyl thio-β-galactoside (IPTG). To enhance translocon delivery into cell membranes, for experiments analyzing IpaB during cellular infection, strains also expressed the *E. coli* adhesion Afa-1 (24).

### Cell culture

HeLa (CCL2) cells (ATCC) were cultured in Dulbecco’s Modified Eagles media (DMEM) supplemented with 0.45% glucose and 10% heat-inactivated fetal bovine serum (FBS) and were maintained at 37°C with 5% CO_2_ and humidity.

### Quantification of intracellular bacteria

MEF cells were seeded at 1.5 × 10^4^ cells/well in 96-well plates the day prior to infection. Exponential phase *S. flexneri* in were added to cell monolayers in Hanks balanced salt solution (HBSS) at an (multiplicity of infection) MOI of 100, were centrifuged onto cells for 10 min at 2,000 rpm. Extracellular bacteria were removed by washing at 20 min of infection, and remaining extracellular but not intracellular bacteria were killed by incubation of the coculture for an additional hour at 37°C with HBSS containing 25 μg/ml gentamicin. Intracellular bacteria were quantified by plating lysates in PBS with 0.5% Triton X-100 on selective media.

### Secretion assay

Secretion via the T3SS was induced with Congo red (8, 25, 26). Briefly, exponential phase cultures, induced for 2 hours with 1mM IPTG, were normalized by OD_600_. Normalized bacteria were incubated stationary at 37°C for 45 minutes in PBS containing 1mM IPTG and 10μM Congo red. Supernatants were analyzed after removal of bacteria by centrifugation.

### Erythrocyte lysis assay

Efficiency of pore formation by *S. flexneri* expressing IpaB derivatives was assayed via quantification of contact dependent sheep erythrocyte hemolysis (5, 8). Briefly, 10^8^ erythrocytes were infected by centrifugation of *S. flexneri* at a MOI of 25 in 30mM Tris pH 7.4 at 25°C for 10 minutes at 2,000 x *g* and incubation at 37°C in 5% CO_2_ for 30 minutes. Co-cultures were mixed by pipetting. Supernatants were isolated by centrifugation at 25°C for 10 minutes at 2,000 x *g*. Hemolysis was quantified spectrophotometrically at A_570_ using a Wallac 1,420 Victor^2^ microplate reader (Perkin Elmer).

### Cysteine labeling with PEG5000-maleimide

Individual IpaB amino acids were replaced with cysteine, and accessibility of IpaB cysteines in soluble IpaB or in membrane-embedded pores was as described for IpaC (11, 18). In brief, soluble IpaB was analyzed by incubating Congo red-induced culture supernatants with 2.5mM PEG5000-maleimide for 30 minutes at 30°C. IpaB in membrane-embedded translocons was analyzed upon infection of HeLa monolayers with bacteria (MOI of 100 in 50mM Tris with 150mM NaCl, 1mM IPTG, and 2.5mM PEG5000-maleimide) by centrifugation for 10 min at 2,000 rpm, harvesting infected cells after 20 minutes of additional incubation, and isolating plasma membrane-enriched fractions, which contains translocons, via sequential membrane fractionation with 0.2% saponin then 0.5% Triton X-100, as described (8, 27).

### Covalent crosslinking of cysteine residues

Crosslinking of IpaB cysteine substitutions with cysteine substitutions in neighboring IpaB molecules was induced using the oxidant copper phenanthroline, as described (10, 28). Briefly, HeLa monolayers were infected with bacteria, at a MOI of 100 in HBSS with 4% FBS, 1mM IPTG, and 25μM copper phenanthroline, with centrifugation (as above), were incubated for 20 minutes longer, then were lysed with 0.5% Triton X-100. The plasma membrane-containing detergent soluble fraction was recovered by centrifugation. Exposure of cells to 25μM copper phenanthroline did not induce cell detachment, suggesting that the cytosolic redox environment was not significantly altered (29).

### Statistical analysis

Except where specifically noted, all data are from three independent experiments and the means ± standard errors of the means (SEM) are presented. Dots within graphs represent independent experimental replicates. The means were compared among groups by a one-way analysis of variance (ANOVA) with appropriate *post hoc* test using GraphPad Prism 8 (GraphPad Software, Inc.). Signal from western blots was captured by film, film was digitized using an Epson Perfection 4990 photo scanner, and the density of bands was determined using ImageJ (National Institutes of Health).

## Acknowledgments

We thank Goldberg laboratory members for critical reading of the manuscript. This work was funded by NIH grants R01 AI081724 (to MBG), F32 AI147549 (to PC), T32 AI007061 (to PC and BCR), and by NIH grant F32 AI114162, the Massachusetts General Hospital Executive Committee on Research Tosteson Award, and the Charles A. King Trust Postdoctoral Research Fellowship Program, Bank of America, N.A., Co-Trustees (to BCR). The funders had no role in study design, data collection and interpretation, or the decision to submit the work for publication. The authors have no financial or other relationships that are relevant to the study.

Study conceptualization was by MBG, PC, and BCR; investigation by PC, BCR, JKD-L, NB, and KE; methodology and formal analysis by PC; visualization by PC and MBG; writing of the original draft by PC; and review and editing by BCR, JKD-L, KE, and MBG.

