## Supplemental data for "Topology and contribution to the pore channel lining of plasma membrane-embedded *S. flexneri* type 3 secretion translocase IpaB"

### Supplemental Figures

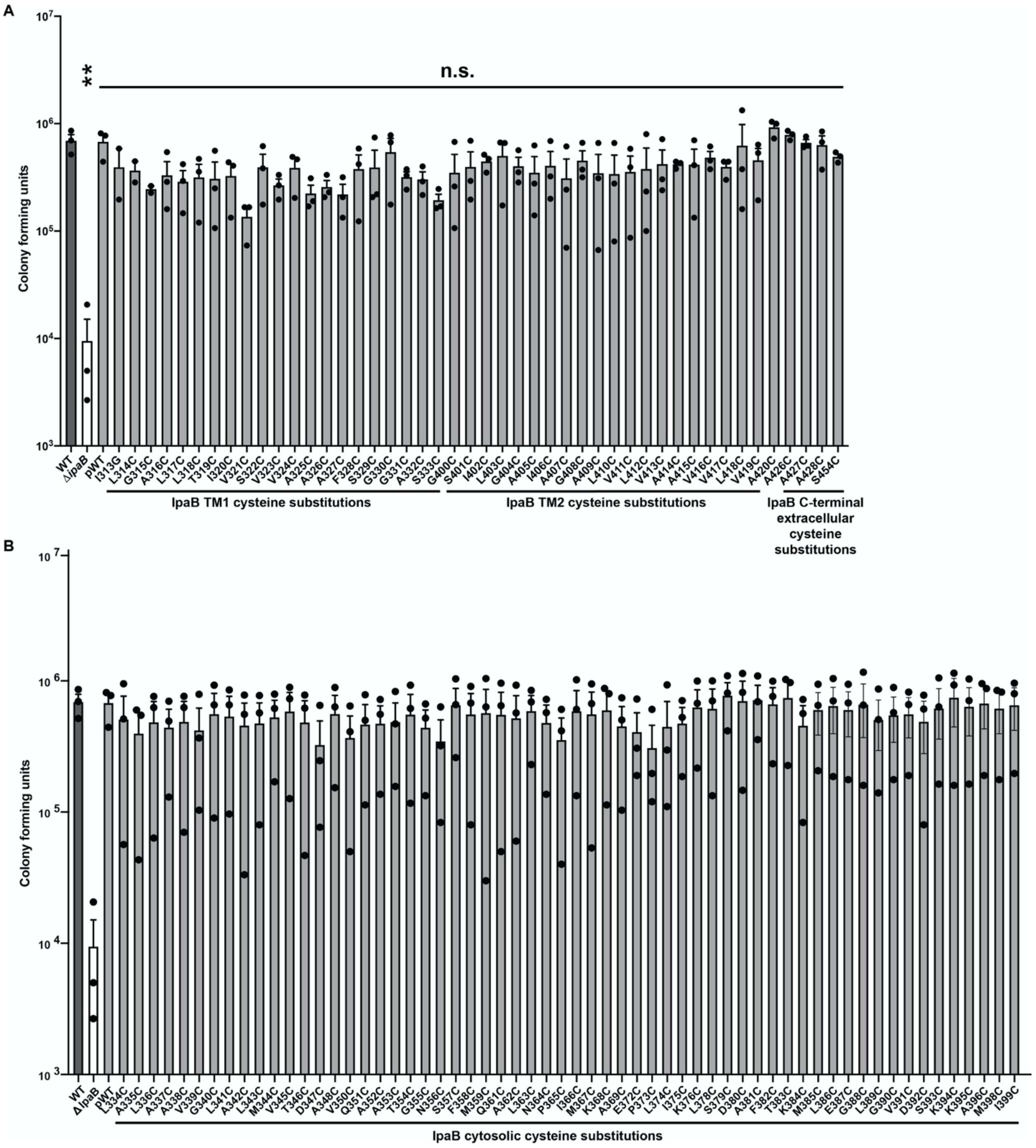

**Supplemental Figure 1.** Cysteine substitutions in IpaB do not alter *S. flexneri* invasion into host cells. Infection of HeLa monolayers with *S. flexneri* ∆*ipaB* expressing wildtype IpaB or a single IpaB cysteine substitution derivative. Positive control, wildtype *S. flexneri*. Negative control, *S. flexneri* ∆*ipaB.* Quantification of colony forming units from three independent experiments. (A) IpaB cysteine derivatives within TM1, TM2, and C-terminal extracellular domain. Means ± SEM are plotted. Black dots represent values obtained from individual experiments. **, $p<0.01$; ANOVA with Dunnett's *post hoc* test comparing colony forming units of *S. flexneri* ∆*ipaB, S. flexneri* ∆*ipaB* expressing each IpaB cysteine derivative, and *S. flexneri* ∆*ipaB* expressing WT IpaB (pWT) to colony forming units of WT *S. flexneri*. (B) IpaB cysteine derivatives within the cytosolic region. The trends observed for data in panel B do not reach statistical significance.

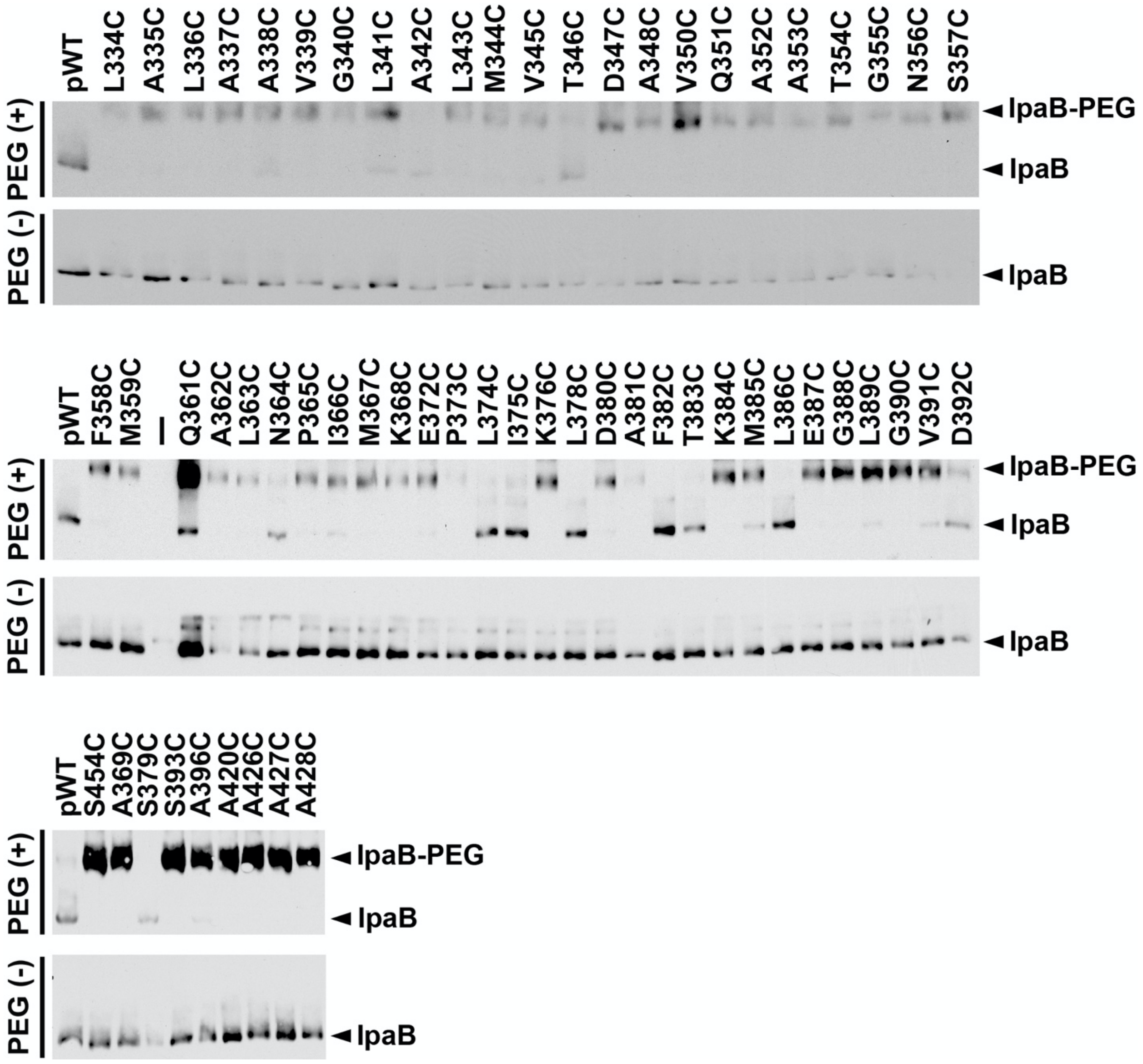


**Supplemental Figure 2.** In the context of soluble IpaB, most cysteine substitutions along the cytosolic domain of IpaB are accessible. Gel migration of PEG5000-maleimide labeled (IpaB-PEG) or unlabeled (IpaB) soluble IpaB in culture supernatants of indicated strains following chemical activation of type 3 secretion with Congo red. *S. flexneri* Δ*ipaB* expressing wildtype (pWT) IpaB or a single IpaB cysteine substitution derivative. Representative western blots.

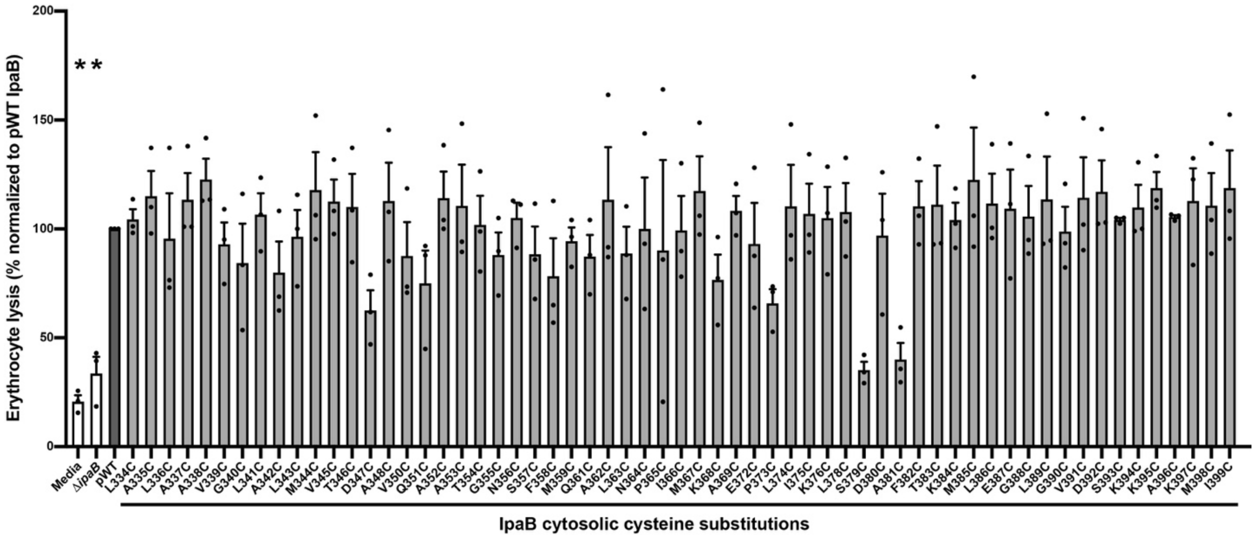


**Supplemental Figure 3.** Efficiency of pore formation by *S. flexneri* strains expressing single cysteine substitutions in IpaB. Quantification of hemoglobin release upon infection of sheep erythrocytes with *S. flexneri ∆ipaB*, *S. flexneri ∆ipaB* expressing wildtype IpaB (pWT) or a single cysteine substitution derivative. The abundance of hemoglobin release was quantified at $A_{570}$ from at least three independent experiments. Means ± SEM are plotted. Black dots represent values obtained from individual experiments. *, p<0.05; ANOVA with Dunnett's *post hoc* test comparing each cysteine substitution mutant to *S. flexneri ∆ipaB* producing WT IpaB (pWT); the difference for each other strain is not significant.

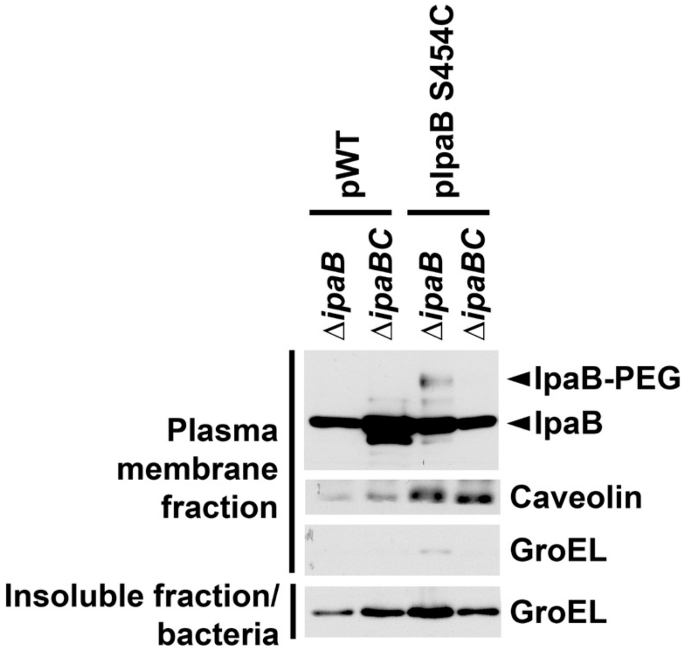


**Supplemental Figure 4.** In the absence of IpaC, membrane-associated IpaB C-terminal cysteine substitution does not react to PEG5000-maleimide. Gel migration of PEG5000-maleimide labeled (IpaB-PEG) or unlabeled (IpaB) IpaB in plasma membrane enriched fractions. *S. flexneri* Δ*ipaB* or Δ*ipaBC* expressing wildtype (pWT) IpaB or the IpaB cysteine substitution derivative S454C (pIpaB S454C). Positive control, *S. flexneri* Δ*ipaB* expressing IpaB S454. IpaB S454C lies in the C-terminal extracellular domain and is accessible to PEG5000-maleimide labeling in membrane inserted translocons (Fig. 3). Caveolin-1, plasma membrane marker; GroEL, bacterial cytosolic protein.

**Supplemental Table**

**Table S1. Strains used in this study**

| **Strain** | **Plasmid 1** | **Plasmid 2** | **Relevant genotype** | **Source or reference** |
|---|---|---|---|---|
| *E. coli* DH10B | | | $F^{-}$ *mcr*A Δ(*mrr-hsd*RMS-*mcr*BC) φ80*lac*ZΔM15 Δ*lac*X74 *rec*A1 *end*A1 *ara*D139 Δ(*ara-leu*)7697 *gal*U *gal*K $\lambda^{-}$ *rps*L($Str^{R}$) *nup*G | Thermo Fisher (18290015) |
| *E. coli* HB101 | pIL22 | | pIL22 is pBR322 containing the pBR322 containing the AFA-I adhesin of uropathogenic E. coli K552 | (1) |
| *S. flexneri* 2457T | | | Wildtype serotype 2a | (2) |
| *S. flexneri* 2457T Δ*ipaB* | | | Deletion of *ipaB* | This study |
| *S. flexneri* 2457T Δ*acp* | | | Deletion of *acp* | This study |
| *S. flexneri* 2457T Δ*ipaB* | pDSW206-wildtype IpaB | | Δ*ipaB* expressing wildtype IpaB | This study |

| | | | | |
|---|---|---|---|---|
| *S. flexneri* 2457T Δ*ipaB* | pDSW206-IpaB C309S | | Δ*ipaB* expressing IpaB C309S | This study |
| *S. flexneri* 2457T Δ*ipaB* | pDSW206-IpaB I313C | | Δ*ipaB* expressing IpaB I313C | This study |
| *S. flexneri* 2457T Δ*ipaB* | pDSW206-IpaB L314C | | Δ*ipaB* expressing IpaB L314C | This study |
| *S. flexneri* 2457T Δ*ipaB* | pDSW206-IpaB G315C | | Δ*ipaB* expressing IpaB G315C | This study |
| *S. flexneri* 2457T Δ*ipaB* | pDSW206-IpaB A316C | | Δ*ipaB* expressing IpaB A316C | This study |
| *S. flexneri* 2457T Δ*ipaB* | pDSW206-IpaB L317C | | Δ*ipaB* expressing IpaB L317C | This study |
| *S. flexneri* 2457T Δ*ipaB* | pDSW206-IpaB L318C | | Δ*ipaB* expressing IpaB L318C | This study |
| *S. flexneri* 2457T Δ*ipaB* | pDSW206-IpaB T319C | | Δ*ipaB* expressing IpaB T319C | This study |
| *S. flexneri* 2457T Δ*ipaB* | pDSW206-IpaB I320C | | Δ*ipaB* expressing IpaB I320C | This study |
| *S. flexneri* 2457T Δ*ipaB* | pDSW206-IpaB V321C | | Δ*ipaB* expressing IpaB V321C | This study |
| *S. flexneri* 2457T Δ*ipaB* | pDSW206-IpaB S322C | | Δ*ipaB* expressing IpaB S322C | This study |
| *S. flexneri* 2457T Δ*ipaB* | pDSW206-IpaB V323C | | Δ*ipaB* expressing IpaB V323C | This study |
| *S. flexneri* 2457T Δ*ipaB* | pDSW206-IpaB V324C | | Δ*ipaB* expressing IpaB V324C | This study |
| *S. flexneri* 2457T Δ*ipaB* | pDSW206-IpaB A325C | | Δ*ipaB* expressing IpaB A325C | This study |
| *S. flexneri* 2457T Δ*ipaB* | pDSW206-IpaB A326C | | Δ*ipaB* expressing IpaB A326C | This study |
| *S. flexneri* 2457T Δ*ipaB* | pDSW206-IpaB A327C | | Δ*ipaB* expressing IpaB A327C | This study |
| *S. flexneri* 2457T Δ*ipaB* | pDSW206-IpaB F328C | | Δ*ipaB* expressing IpaB F328C | This study |

| | | | | |
|---|---|---|---|---|
| *S. flexneri* 2457T Δ*ipaB* | pDSW206-IpaB S329C | | Δ*ipaB* expressing IpaB S329C | This study |
| *S. flexneri* 2457T Δ*ipaB* | pDSW206-IpaB G330C | | Δ*ipaB* expressing IpaB G330C | This study |
| *S. flexneri* 2457T Δ*ipaB* | pDSW206-IpaB G331C | | Δ*ipaB* expressing IpaB G331C | This study |
| *S. flexneri* 2457T Δ*ipaB* | pDSW206-IpaB A332C | | Δ*ipaB* expressing IpaB A332C | This study |
| *S. flexneri* 2457T Δ*ipaB* | pDSW206-IpaB S333C | | Δ*ipaB* expressing IpaB S333C | This study |
| *S. flexneri* 2457T Δ*ipaB* | pDSW206-IpaB L334C | | Δ*ipaB* expressing IpaB L334C | This study |
| *S. flexneri* 2457T Δ*ipaB* | pDSW206-IpaB A335C | | Δ*ipaB* expressing IpaB A335C | This study |
| *S. flexneri* 2457T Δ*ipaB* | pDSW206-IpaB L336C | | Δ*ipaB* expressing IpaB L336C | This study |
| *S. flexneri* 2457T Δ*ipaB* | pDSW206-IpaB A337C | | Δ*ipaB* expressing IpaB A337C | This study |
| *S. flexneri* 2457T Δ*ipaB* | pDSW206-IpaB A338C | | Δ*ipaB* expressing IpaB A338C | This study |
| *S. flexneri* 2457T Δ*ipaB* | pDSW206-IpaB V339C | | Δ*ipaB* expressing IpaB V339C | This study |
| *S. flexneri* 2457T Δ*ipaB* | pDSW206-IpaB G340C | | Δ*ipaB* expressing IpaB G340C | This study |
| *S. flexneri* 2457T Δ*ipaB* | pDSW206-IpaB L341C | | Δ*ipaB* expressing IpaB L341C | This study |
| *S. flexneri* 2457T Δ*ipaB* | pDSW206-IpaB A342C | | Δ*ipaB* expressing IpaB A342C | This study |
| *S. flexneri* 2457T Δ*ipaB* | pDSW206-IpaB L343C | | Δ*ipaB* expressing IpaB L343C | This study |
| *S. flexneri* 2457T Δ*ipaB* | pDSW206-IpaB M344C | | Δ*ipaB* expressing IpaB M344C | This study |
| *S. flexneri* 2457T Δ*ipaB* | pDSW206-IpaB V345C | | Δ*ipaB* expressing IpaB V345C | This study |

| | | | | |
|---|---|---|---|---|
| *S. flexneri* 2457T Δ*ipaB* | pDSW206-IpaB T346C | | Δ*ipaB* expressing IpaB T346C | This study |
| *S. flexneri* 2457T Δ*ipaB* | pDSW206-IpaB D347C | | Δ*ipaB* expressing IpaB D347C | This study |
| *S. flexneri* 2457T Δ*ipaB* | pDSW206-IpaB A348C | | Δ*ipaB* expressing IpaB A348C | This study |
| *S. flexneri* 2457T Δ*ipaB* | pDSW206-IpaB V350C | | Δ*ipaB* expressing IpaB V350C | This study |
| *S. flexneri* 2457T Δ*ipaB* | pDSW206-IpaB Q351C | | Δ*ipaB* expressing IpaB Q351C | This study |
| *S. flexneri* 2457T Δ*ipaB* | pDSW206-IpaB A352C | | Δ*ipaB* expressing IpaB A352C | This study |
| *S. flexneri* 2457T Δ*ipaB* | pDSW206-IpaB A353C | | Δ*ipaB* expressing IpaB A353C | This study |
| *S. flexneri* 2457T Δ*ipaB* | pDSW206-IpaB T354C | | Δ*ipaB* expressing IpaB T354C | This study |
| *S. flexneri* 2457T Δ*ipaB* | pDSW206-IpaB G355C | | Δ*ipaB* expressing IpaB G355C | This study |
| *S. flexneri* 2457T Δ*ipaB* | pDSW206-IpaB N356C | | Δ*ipaB* expressing IpaB N356C | This study |
| *S. flexneri* 2457T Δ*ipaB* | pDSW206-IpaB S357C | | Δ*ipaB* expressing IpaB S357C | This study |
| *S. flexneri* 2457T Δ*ipaB* | pDSW206-IpaB F358C | | Δ*ipaB* expressing IpaB F358C | This study |
| *S. flexneri* 2457T Δ*ipaB* | pDSW206-IpaB M359C | | Δ*ipaB* expressing IpaB M359C | This study |
| *S. flexneri* 2457T Δ*ipaB* | pDSW206-IpaB Q361C | | Δ*ipaB* expressing IpaB Q361C | This study |
| *S. flexneri* 2457T Δ*ipaB* | pDSW206-IpaB A362C | | Δ*ipaB* expressing IpaB A362C | This study |
| *S. flexneri* 2457T Δ*ipaB* | pDSW206-IpaB L363C | | Δ*ipaB* expressing IpaB L363C | This study |
| *S. flexneri* 2457T Δ*ipaB* | pDSW206-IpaB N364C | | Δ*ipaB* expressing IpaB N364C | This study |

| | | | | |
|---|---|---|---|---|
| *S. flexneri* 2457T Δ*ipaB* | pDSW206-IpaB P365C | | Δ*ipaB* expressing IpaB P365C | This study |
| *S. flexneri* 2457T Δ*ipaB* | pDSW206-IpaB I366C | | Δ*ipaB* expressing IpaB I366C | This study |
| *S. flexneri* 2457T Δ*ipaB* | pDSW206-IpaB M367C | | Δ*ipaB* expressing IpaB M367C | This study |
| *S. flexneri* 2457T Δ*ipaB* | pDSW206-IpaB K368C | | Δ*ipaB* expressing IpaB K368C | This study |
| *S. flexneri* 2457T Δ*ipaB* | pDSW206-IpaB A369C | | Δ*ipaB* expressing IpaB A369C | This study |
| *S. flexneri* 2457T Δ*ipaB* | pDSW206-IpaB E372C | | Δ*ipaB* expressing IpaB E372C | This study |
| *S. flexneri* 2457T Δ*ipaB* | pDSW206-IpaB P373C | | Δ*ipaB* expressing IpaB P373C | This study |
| *S. flexneri* 2457T Δ*ipaB* | pDSW206-IpaB L374C | | Δ*ipaB* expressing IpaB L374C | This study |
| *S. flexneri* 2457T Δ*ipaB* | pDSW206-IpaB I375C | | Δ*ipaB* expressing IpaB I375C | This study |
| *S. flexneri* 2457T Δ*ipaB* | pDSW206-IpaB K376C | | Δ*ipaB* expressing IpaB K376C | This study |
| *S. flexneri* 2457T Δ*ipaB* | pDSW206-IpaB L378C | | Δ*ipaB* expressing IpaB L378C | This study |
| *S. flexneri* 2457T Δ*ipaB* | pDSW206-IpaB S379C | | Δ*ipaB* expressing IpaB S379C | This study |
| *S. flexneri* 2457T Δ*ipaB* | pDSW206-IpaB D380C | | Δ*ipaB* expressing IpaB D380C | This study |
| *S. flexneri* 2457T Δ*ipaB* | pDSW206-IpaB A381C | | Δ*ipaB* expressing IpaB A381C | This study |
| *S. flexneri* 2457T Δ*ipaB* | pDSW206-IpaB F382C | | Δ*ipaB* expressing IpaB F382C | This study |
| *S. flexneri* 2457T Δ*ipaB* | pDSW206-IpaB T383C | | Δ*ipaB* expressing IpaB T383C | This study |
| *S. flexneri* 2457T Δ*ipaB* | pDSW206-IpaB K384C | | Δ*ipaB* expressing IpaB K384C | This study |

| | | | | |
|---|---|---|---|---|
| *S. flexneri* 2457T Δ*ipaB* | pDSW206-IpaB M385C | | Δ*ipaB* expressing IpaB M385C | This study |
| *S. flexneri* 2457T Δ*ipaB* | pDSW206-IpaB L386C | | Δ*ipaB* expressing IpaB L386C | This study |
| *S. flexneri* 2457T Δ*ipaB* | pDSW206-IpaB E387C | | Δ*ipaB* expressing IpaB E387C | This study |
| *S. flexneri* 2457T Δ*ipaB* | pDSW206-IpaB G388C | | Δ*ipaB* expressing IpaB G388C | This study |
| *S. flexneri* 2457T Δ*ipaB* | pDSW206-IpaB L389C | | Δ*ipaB* expressing IpaB L389C | This study |
| *S. flexneri* 2457T Δ*ipaB* | pDSW206-IpaB G390C | | Δ*ipaB* expressing IpaB G390C | This study |
| *S. flexneri* 2457T Δ*ipaB* | pDSW206-IpaB V391C | | Δ*ipaB* expressing IpaB V391C | This study |
| *S. flexneri* 2457T Δ*ipaB* | pDSW206-IpaB D392C | | Δ*ipaB* expressing IpaB D392C | This study |
| *S. flexneri* 2457T Δ*ipaB* | pDSW206-IpaB S393C | | Δ*ipaB* expressing IpaB S393C | This study |
| *S. flexneri* 2457T Δ*ipaB* | pDSW206-IpaB K394C | | Δ*ipaB* expressing IpaB K394C | This study |
| *S. flexneri* 2457T Δ*ipaB* | pDSW206-IpaB K395C | | Δ*ipaB* expressing IpaB K395C | This study |
| *S. flexneri* 2457T Δ*ipaB* | pDSW206-IpaB A396C | | Δ*ipaB* expressing IpaB A396C | This study |
| *S. flexneri* 2457T Δ*ipaB* | pDSW206-IpaB M398C | | Δ*ipaB* expressing IpaB M398C | This study |
| *S. flexneri* 2457T Δ*ipaB* | pDSW206-IpaB I399C | | Δ*ipaB* expressing IpaB I399C | This study |
| *S. flexneri* 2457T Δ*ipaB* | pDSW206-IpaB G400C | | Δ*ipaB* expressing IpaB G400C | This study |
| *S. flexneri* 2457T Δ*ipaB* | pDSW206-IpaB S401C | | Δ*ipaB* expressing IpaB S401C | This study |
| *S. flexneri* 2457T Δ*ipaB* | pDSW206-IpaB I402C | | Δ*ipaB* expressing IpaB I402C | This study |

| | | | | |
|---|---|---|---|---|
| *S. flexneri* 2457T Δ*ipaB* | pDSW206-IpaB L403C | | Δ*ipaB* expressing IpaB L403C | This study |
| *S. flexneri* 2457T Δ*ipaB* | pDSW206-IpaB G404C | | Δ*ipaB* expressing IpaB G404C | This study |
| *S. flexneri* 2457T Δ*ipaB* | pDSW206-IpaB A405C | | Δ*ipaB* expressing IpaB A405C | This study |
| *S. flexneri* 2457T Δ*ipaB* | pDSW206-IpaB I406C | | Δ*ipaB* expressing IpaB I406C | This study |
| *S. flexneri* 2457T Δ*ipaB* | pDSW206-IpaB A407C | | Δ*ipaB* expressing IpaB A407C | This study |
| *S. flexneri* 2457T Δ*ipaB* | pDSW206-IpaB G408C | | Δ*ipaB* expressing IpaB G408C | This study |
| *S. flexneri* 2457T Δ*ipaB* | pDSW206-IpaB A409C | | Δ*ipaB* expressing IpaB A409C | This study |
| *S. flexneri* 2457T Δ*ipaB* | pDSW206-IpaB L410C | | Δ*ipaB* expressing IpaB L410C | This study |
| *S. flexneri* 2457T Δ*ipaB* | pDSW206-IpaB V411C | | Δ*ipaB* expressing IpaB V411C | This study |
| *S. flexneri* 2457T Δ*ipaB* | pDSW206-IpaB L412C | | Δ*ipaB* expressing IpaB L412C | This study |
| *S. flexneri* 2457T Δ*ipaB* | pDSW206-IpaB V413C | | Δ*ipaB* expressing IpaB V413C | This study |
| *S. flexneri* 2457T Δ*ipaB* | pDSW206-IpaB A414C | | Δ*ipaB* expressing IpaB A414C | This study |
| *S. flexneri* 2457T Δ*ipaB* | pDSW206-IpaB A415C | | Δ*ipaB* expressing IpaB A415C | This study |
| *S. flexneri* 2457T Δ*ipaB* | pDSW206-IpaB V416C | | Δ*ipaB* expressing IpaB V416C | This study |
| *S. flexneri* 2457T Δ*ipaB* | pDSW206-IpaB V417C | | Δ*ipaB* expressing IpaB V417C | This study |
| *S. flexneri* 2457T Δ*ipaB* | pDSW206-IpaB L418C | | Δ*ipaB* expressing IpaB L418C | This study |
| *S. flexneri* 2457T Δ*ipaB* | pDSW206-IpaB V419C | | Δ*ipaB* expressing IpaB V419C | This study |

| | | | | |
|---|---|---|---|---|
| *S. flexneri* 2457T Δ*ipaB* | pDSW206-IpaB A420C | | Δ*ipaB* expressing IpaB A420C | This study |
| *S. flexneri* 2457T Δ*ipaB* | pDSW206-IpaB A426C | | Δ*ipaB* expressing IpaB A426C | This study |
| *S. flexneri* 2457T Δ*ipaB* | pDSW206-IpaB A427C | | Δ*ipaB* expressing IpaB A427C | This study |
| *S. flexneri* 2457T Δ*ipaB* | pDSW206-IpaB A428C | | Δ*ipaB* expressing IpaB A428C | This study |
| *S. flexneri* 2457T Δ*ipaB* | pDSW206-IpaB S454C | | Δ*ipaB* expressing IpaB S454C | This study |
| *S. flexneri* 2457T Δ*ipaB* | pDSW206-wildtype IpaB | pNG162-Afa1 | Δ*ipaB* expressing wildtype IpaB and adhesin | Laboratory stock |
| *S. flexneri* 2457T Δ*ipaB* | pDSW206-IpaB I313C | pNG162-Afa1 | Δ*ipaB* expressing IpaB I313C and adhesin | This study |
| *S. flexneri* 2457T Δ*ipaB* | pDSW206-IpaB L314C | pNG162-Afa1 | Δ*ipaB* expressing IpaB L314C and adhesin | This study |
| *S. flexneri* 2457T Δ*ipaB* | pDSW206-IpaB G315C | pNG162-Afa1 | Δ*ipaB* expressing IpaB G315C and adhesin | This study |
| *S. flexneri* 2457T Δ*ipaB* | pDSW206-IpaB A316C | pNG162-Afa1 | Δ*ipaB* expressing IpaB A316C and adhesin | This study |
| *S. flexneri* 2457T Δ*ipaB* | pDSW206-IpaB L317C | pNG162-Afa1 | Δ*ipaB* expressing IpaB L317C and adhesin | This study |

| | | | | |
|---|---|---|---|---|
| *S. flexneri* 2457T Δ*ipaB* | pDSW206-IpaB L318C | pNG162-Afa1 | Δ*ipaB* expressing IpaB L318C and adhesin | This study |
| *S. flexneri* 2457T Δ*ipaB* | pDSW206-IpaB T319C | pNG162-Afa1 | Δ*ipaB* expressing IpaB T319C and adhesin | This study |
| *S. flexneri* 2457T Δ*ipaB* | pDSW206-IpaB I320C | pNG162-Afa1 | Δ*ipaB* expressing IpaB I320C and adhesin | This study |
| *S. flexneri* 2457T Δ*ipaB* | pDSW206-IpaB V321C | pNG162-Afa1 | Δ*ipaB* expressing IpaB V321C and adhesin | This study |
| *S. flexneri* 2457T Δ*ipaB* | pDSW206-IpaB S322C | pNG162-Afa1 | Δ*ipaB* expressing IpaB S322C and adhesin | This study |
| *S. flexneri* 2457T Δ*ipaB* | pDSW206-IpaB V323C | pNG162-Afa1 | Δ*ipaB* expressing IpaB V323C and adhesin | This study |
| *S. flexneri* 2457T Δ*ipaB* | pDSW206-IpaB V324C | pNG162-Afa1 | Δ*ipaB* expressing IpaB V324C and adhesin | This study |
| *S. flexneri* 2457T Δ*ipaB* | pDSW206-IpaB A325C | pNG162-Afa1 | Δ*ipaB* expressing IpaB A325C and adhesin | This study |

| | | | | |
|---|---|---|---|---|
| *S. flexneri* 2457T Δ*ipaB* | pDSW206-IpaB A326C | pNG162-Afa1 | Δ*ipaB* expressing IpaB A326C and adhesin | This study |
| *S. flexneri* 2457T Δ*ipaB* | pDSW206-IpaB A327C | pNG162-Afa1 | Δ*ipaB* expressing IpaB A327C and adhesin | This study |
| *S. flexneri* 2457T Δ*ipaB* | pDSW206-IpaB F328C | pNG162-Afa1 | Δ*ipaB* expressing IpaB F328C and adhesin | This study |
| *S. flexneri* 2457T Δ*ipaB* | pDSW206-IpaB S329C | pNG162-Afa1 | Δ*ipaB* expressing IpaB S329C and adhesin | This study |
| *S. flexneri* 2457T Δ*ipaB* | pDSW206-IpaB G330C | pNG162-Afa1 | Δ*ipaB* expressing IpaB G330C and adhesin | This study |
| *S. flexneri* 2457T Δ*ipaB* | pDSW206-IpaB G331C | pNG162-Afa1 | Δ*ipaB* expressing IpaB G331C and adhesin | This study |
| *S. flexneri* 2457T Δ*ipaB* | pDSW206-IpaB A332C | pNG162-Afa1 | Δ*ipaB* expressing IpaB A332C and adhesin | This study |
| *S. flexneri* 2457T Δ*ipaB* | pDSW206-IpaB S333C | pNG162-Afa1 | Δ*ipaB* expressing IpaB S333C and adhesin | This study |

| | | | | |
|---|---|---|---|---|
| *S. flexneri* 2457T Δ*ipaB* | pDSW206-IpaB L334C | pNG162-Afa1 | Δ*ipaB* expressing IpaB L334C and adhesin | This study |
| *S. flexneri* 2457T Δ*ipaB* | pDSW206-IpaB A335C | pNG162-Afa1 | Δ*ipaB* expressing IpaB A335C and adhesin | This study |
| *S. flexneri* 2457T Δ*ipaB* | pDSW206-IpaB L336C | pNG162-Afa1 | Δ*ipaB* expressing IpaB L336C and adhesin | This study |
| *S. flexneri* 2457T Δ*ipaB* | pDSW206-IpaB A337C | pNG162-Afa1 | Δ*ipaB* expressing IpaB A337C and adhesin | This study |
| *S. flexneri* 2457T Δ*ipaB* | pDSW206-IpaB A338C | pNG162-Afa1 | Δ*ipaB* expressing IpaB A338C and adhesin | This study |
| *S. flexneri* 2457T Δ*ipaB* | pDSW206-IpaB V339C | pNG162-Afa1 | Δ*ipaB* expressing IpaB V339C and adhesin | This study |
| *S. flexneri* 2457T Δ*ipaB* | pDSW206-IpaB G340C | pNG162-Afa1 | Δ*ipaB* expressing IpaB G340C and adhesin | This study |
| *S. flexneri* 2457T Δ*ipaB* | pDSW206-IpaB L341C | pNG162-Afa1 | Δ*ipaB* expressing IpaB L341C and adhesin | This study |

| | | | | |
|---|---|---|---|---|
| *S. flexneri* 2457T Δ*ipaB* | pDSW206-IpaB A342C | pNG162-Afa1 | Δ*ipaB* expressing IpaB A342C and adhesin | This study |
| *S. flexneri* 2457T Δ*ipaB* | pDSW206-IpaB L343C | pNG162-Afa1 | Δ*ipaB* expressing IpaB L343C and adhesin | This study |
| *S. flexneri* 2457T Δ*ipaB* | pDSW206-IpaB M344C | pNG162-Afa1 | Δ*ipaB* expressing IpaB M344C and adhesin | This study |
| *S. flexneri* 2457T Δ*ipaB* | pDSW206-IpaB V345C | pNG162-Afa1 | Δ*ipaB* expressing IpaB V345C and adhesin | This study |
| *S. flexneri* 2457T Δ*ipaB* | pDSW206-IpaB T346C | pNG162-Afa1 | Δ*ipaB* expressing IpaB T346C and adhesin | This study |
| *S. flexneri* 2457T Δ*ipaB* | pDSW206-IpaB D347C | pNG162-Afa1 | Δ*ipaB* expressing IpaB D347C and adhesin | This study |
| *S. flexneri* 2457T Δ*ipaB* | pDSW206-IpaB A348C | pNG162-Afa1 | Δ*ipaB* expressing IpaB A348C and adhesin | This study |
| *S. flexneri* 2457T Δ*ipaB* | pDSW206-IpaB V350C | pNG162-Afa1 | Δ*ipaB* expressing IpaB V350C and adhesin | This study |

| | | | | |
|---|---|---|---|---|
| *S. flexneri* 2457T Δ*ipaB* | pDSW206-IpaB Q351C | pNG162-Afa1 | Δ*ipaB* expressing IpaB Q351C and adhesin | This study |
| *S. flexneri* 2457T Δ*ipaB* | pDSW206-IpaB A352C | pNG162-Afa1 | Δ*ipaB* expressing IpaB A352C and adhesin | This study |
| *S. flexneri* 2457T Δ*ipaB* | pDSW206-IpaB A353C | pNG162-Afa1 | Δ*ipaB* expressing IpaB A353C and adhesin | This study |
| *S. flexneri* 2457T Δ*ipaB* | pDSW206-IpaB T354C | pNG162-Afa1 | Δ*ipaB* expressing IpaB T354C and adhesin | This study |
| *S. flexneri* 2457T Δ*ipaB* | pDSW206-IpaB G355C | pNG162-Afa1 | Δ*ipaB* expressing IpaB G355C and adhesin | This study |
| *S. flexneri* 2457T Δ*ipaB* | pDSW206-IpaB N356C | pNG162-Afa1 | Δ*ipaB* expressing IpaB N356C and adhesin | This study |
| *S. flexneri* 2457T Δ*ipaB* | pDSW206-IpaB S357C | pNG162-Afa1 | Δ*ipaB* expressing IpaB S357C and adhesin | This study |
| *S. flexneri* 2457T Δ*ipaB* | pDSW206-IpaB F358C | pNG162-Afa1 | Δ*ipaB* expressing IpaB F358C and adhesin | This study |

| | | | | |
|---|---|---|---|---|
| *S. flexneri* 2457T Δ*ipaB* | pDSW206-IpaB M359C | pNG162-Afa1 | Δ*ipaB* expressing IpaB M359C and adhesin | This study |
| *S. flexneri* 2457T Δ*ipaB* | pDSW206-IpaB Q361C | pNG162-Afa1 | Δ*ipaB* expressing IpaB Q361C and adhesin | This study |
| *S. flexneri* 2457T Δ*ipaB* | pDSW206-IpaB A362C | pNG162-Afa1 | Δ*ipaB* expressing IpaB A362C and adhesin | This study |
| *S. flexneri* 2457T Δ*ipaB* | pDSW206-IpaB L363C | pNG162-Afa1 | Δ*ipaB* expressing IpaB L363C and adhesin | This study |
| *S. flexneri* 2457T Δ*ipaB* | pDSW206-IpaB N364C | pNG162-Afa1 | Δ*ipaB* expressing IpaB N364C and adhesin | This study |
| *S. flexneri* 2457T Δ*ipaB* | pDSW206-IpaB P365C | pNG162-Afa1 | Δ*ipaB* expressing IpaB P365C and adhesin | This study |
| *S. flexneri* 2457T Δ*ipaB* | pDSW206-IpaB I366C | pNG162-Afa1 | Δ*ipaB* expressing IpaB I366C and adhesin | This study |
| *S. flexneri* 2457T Δ*ipaB* | pDSW206-IpaB M367C | pNG162-Afa1 | Δ*ipaB* expressing IpaB M367C and adhesin | This study |

| | | | | |
|---|---|---|---|---|
| *S. flexneri* 2457T Δ*ipaB* | pDSW206-IpaB K368C | pNG162-Afa1 | Δ*ipaB* expressing IpaB K368C and adhesin | This study |
| *S. flexneri* 2457T Δ*ipaB* | pDSW206-IpaB A369C | pNG162-Afa1 | Δ*ipaB* expressing IpaB A369C and adhesin | This study |
| *S. flexneri* 2457T Δ*ipaB* | pDSW206-IpaB E372C | pNG162-Afa1 | Δ*ipaB* expressing IpaB E372C and adhesin | This study |
| *S. flexneri* 2457T Δ*ipaB* | pDSW206-IpaB P373C | pNG162-Afa1 | Δ*ipaB* expressing IpaB P373C and adhesin | This study |
| *S. flexneri* 2457T Δ*ipaB* | pDSW206-IpaB L374C | pNG162-Afa1 | Δ*ipaB* expressing IpaB L374C and adhesin | This study |
| *S. flexneri* 2457T Δ*ipaB* | pDSW206-IpaB I375C | pNG162-Afa1 | Δ*ipaB* expressing IpaB I375C and adhesin | This study |
| *S. flexneri* 2457T Δ*ipaB* | pDSW206-IpaB K376C | pNG162-Afa1 | Δ*ipaB* expressing IpaB K376C and adhesin | This study |
| *S. flexneri* 2457T Δ*ipaB* | pDSW206-IpaB L378C | pNG162-Afa1 | Δ*ipaB* expressing IpaB L378C and adhesin | This study |

| | | | | |
|---|---|---|---|---|
| *S. flexneri* 2457T Δ*ipaB* | pDSW206-IpaB S379C | pNG162-Afa1 | Δ*ipaB* expressing IpaB S379C and adhesin | This study |
| *S. flexneri* 2457T Δ*ipaB* | pDSW206-IpaB D380C | pNG162-Afa1 | Δ*ipaB* expressing IpaB D380C and adhesin | This study |
| *S. flexneri* 2457T Δ*ipaB* | pDSW206-IpaB A381C | pNG162-Afa1 | Δ*ipaB* expressing IpaB A381C and adhesin | This study |
| *S. flexneri* 2457T Δ*ipaB* | pDSW206-IpaB F382C | pNG162-Afa1 | Δ*ipaB* expressing IpaB F382C and adhesin | This study |
| *S. flexneri* 2457T Δ*ipaB* | pDSW206-IpaB T383C | pNG162-Afa1 | Δ*ipaB* expressing IpaB T383C and adhesin | This study |
| *S. flexneri* 2457T Δ*ipaB* | pDSW206-IpaB K384C | pNG162-Afa1 | Δ*ipaB* expressing IpaB K384C and adhesin | This study |
| *S. flexneri* 2457T Δ*ipaB* | pDSW206-IpaB M385C | pNG162-Afa1 | Δ*ipaB* expressing IpaB M385C and adhesin | This study |
| *S. flexneri* 2457T Δ*ipaB* | pDSW206-IpaB L386C | pNG162-Afa1 | Δ*ipaB* expressing IpaB L386C and adhesin | This study |

| | | | | |
|---|---|---|---|---|
| *S. flexneri* 2457T Δ*ipaB* | pDSW206-IpaB E387C | pNG162-Afa1 | Δ*ipaB* expressing IpaB E387C and adhesin | This study |
| *S. flexneri* 2457T Δ*ipaB* | pDSW206-IpaB G388C | pNG162-Afa1 | Δ*ipaB* expressing IpaB G388C and adhesin | This study |
| *S. flexneri* 2457T Δ*ipaB* | pDSW206-IpaB L389C | pNG162-Afa1 | Δ*ipaB* expressing IpaB L389C and adhesin | This study |
| *S. flexneri* 2457T Δ*ipaB* | pDSW206-IpaB G390C | pNG162-Afa1 | Δ*ipaB* expressing IpaB G390C and adhesin | This study |
| *S. flexneri* 2457T Δ*ipaB* | pDSW206-IpaB V391C | pNG162-Afa1 | Δ*ipaB* expressing IpaB V391C and adhesin | This study |
| *S. flexneri* 2457T Δ*ipaB* | pDSW206-IpaB D392C | pNG162-Afa1 | Δ*ipaB* expressing IpaB D392C and adhesin | This study |
| *S. flexneri* 2457T Δ*ipaB* | pDSW206-IpaB S393C | pNG162-Afa1 | Δ*ipaB* expressing IpaB S393C and adhesin | This study |
| *S. flexneri* 2457T Δ*ipaB* | pDSW206-IpaB K394C | pNG162-Afa1 | Δ*ipaB* expressing IpaB K394C and adhesin | This study |

| | | | | |
|---|---|---|---|---|
| *S. flexneri* 2457T Δ*ipaB* | pDSW206-IpaB K395C | pNG162-Afa1 | Δ*ipaB* expressing IpaB K395C and adhesin | This study |
| *S. flexneri* 2457T Δ*ipaB* | pDSW206-IpaB A396C | pNG162-Afa1 | Δ*ipaB* expressing IpaB A396C and adhesin | This study |
| *S. flexneri* 2457T Δ*ipaB* | pDSW206-IpaB M398C | pNG162-Afa1 | Δ*ipaB* expressing IpaB M398C and adhesin | This study |
| *S. flexneri* 2457T Δ*ipaB* | pDSW206-IpaB I399C | pNG162-Afa1 | Δ*ipaB* expressing IpaB I399C and adhesin | This study |
| *S. flexneri* 2457T Δ*ipaB* | pDSW206-IpaB G400C | pNG162-Afa1 | Δ*ipaB* expressing IpaB G400C and adhesin | This study |
| *S. flexneri* 2457T Δ*ipaB* | pDSW206-IpaB S401C | pNG162-Afa1 | Δ*ipaB* expressing IpaB S401C and adhesin | This study |
| *S. flexneri* 2457T Δ*ipaB* | pDSW206-IpaB I402C | pNG162-Afa1 | Δ*ipaB* expressing IpaB I402C and adhesin | This study |
| *S. flexneri* 2457T Δ*ipaB* | pDSW206-IpaB L403C | pNG162-Afa1 | Δ*ipaB* expressing IpaB L403C and adhesin | This study |

| | | | | |
|---|---|---|---|---|
| *S. flexneri* 2457T Δ*ipaB* | pDSW206-IpaB G404C | pNG162-Afa1 | Δ*ipaB* expressing IpaB G404C and adhesin | This study |
| *S. flexneri* 2457T Δ*ipaB* | pDSW206-IpaB A405C | pNG162-Afa1 | Δ*ipaB* expressing IpaB A405C and adhesin | This study |
| *S. flexneri* 2457T Δ*ipaB* | pDSW206-IpaB I406C | pNG162-Afa1 | Δ*ipaB* expressing IpaB I406C and adhesin | This study |
| *S. flexneri* 2457T Δ*ipaB* | pDSW206-IpaB A407C | pNG162-Afa1 | Δ*ipaB* expressing IpaB A407C and adhesin | This study |
| *S. flexneri* 2457T Δ*ipaB* | pDSW206-IpaB G408C | pNG162-Afa1 | Δ*ipaB* expressing IpaB G408C and adhesin | This study |
| *S. flexneri* 2457T Δ*ipaB* | pDSW206-IpaB A409C | pNG162-Afa1 | Δ*ipaB* expressing IpaB A409C and adhesin | This study |
| *S. flexneri* 2457T Δ*ipaB* | pDSW206-IpaB L410C | pNG162-Afa1 | Δ*ipaB* expressing IpaB L410C and adhesin | This study |
| *S. flexneri* 2457T Δ*ipaB* | pDSW206-IpaB V411C | pNG162-Afa1 | Δ*ipaB* expressing IpaB V411C and adhesin | This study |

| | | | | |
|---|---|---|---|---|
| *S. flexneri* 2457T Δ*ipaB* | pDSW206-IpaB L412C | pNG162-Afa1 | Δ*ipaB* expressing IpaB L412C and adhesin | This study |
| *S. flexneri* 2457T Δ*ipaB* | pDSW206-IpaB V413C | pNG162-Afa1 | Δ*ipaB* expressing IpaB V413C and adhesin | This study |
| *S. flexneri* 2457T Δ*ipaB* | pDSW206-IpaB A414C | pNG162-Afa1 | Δ*ipaB* expressing IpaB A414C and adhesin | This study |
| *S. flexneri* 2457T Δ*ipaB* | pDSW206-IpaB A415C | pNG162-Afa1 | Δ*ipaB* expressing IpaB A415C and adhesin | This study |
| *S. flexneri* 2457T Δ*ipaB* | pDSW206-IpaB V416C | pNG162-Afa1 | Δ*ipaB* expressing IpaB V416C and adhesin | This study |
| *S. flexneri* 2457T Δ*ipaB* | pDSW206-IpaB V417C | pNG162-Afa1 | Δ*ipaB* expressing IpaB V417C and adhesin | This study |
| *S. flexneri* 2457T Δ*ipaB* | pDSW206-IpaB L418C | pNG162-Afa1 | Δ*ipaB* expressing IpaB L418C and adhesin | This study |
| *S. flexneri* 2457T Δ*ipaB* | pDSW206-IpaB V419C | pNG162-Afa1 | Δ*ipaB* expressing IpaB V419C and adhesin | This study |

| | | | | |
|---|---|---|---|---|
| *S. flexneri* 2457T Δ*ipaB* | pDSW206-IpaB A420C | pNG162-Afa1 | Δ*ipaB* expressing IpaB A420C and adhesin | This study |
| *S. flexneri* 2457T Δ*ipaB* | pDSW206-IpaB A426C | pNG162-Afa1 | Δ*ipaB* expressing IpaB A426C and adhesin | This study |
| *S. flexneri* 2457T Δ*ipaB* | pDSW206-IpaB A427C | pNG162-Afa1 | Δ*ipaB* expressing IpaB A427C and adhesin | This study |
| *S. flexneri* 2457T Δ*ipaB* | pDSW206-IpaB A428C | pNG162-Afa1 | Δ*ipaB* expressing IpaB A428C and adhesin | This study |
| *S. flexneri* 2457T Δ*ipaB* | pDSW206-IpaB S454C | pNG162-Afa1 | Δ*ipaB* expressing IpaB S454C and adhesin | This study |
| *S. flexneri* 2457T Δ*ipaBC* | pDSW206-wildtype IpaB | | Δ*ipaBC* expressing wildtype IpaB | This study |
| *S. flexneri* 2457T Δ*ipaBC* | pDSW206-IpaB S454C | | Δ*ipaBC* expressing IpaB S454C | This study |
